# Hindbrain catecholaminergic inputs to the paraventricular thalamus scale feeding and metabolic efficiency in stress-related contexts

**DOI:** 10.1101/2022.02.03.478953

**Authors:** Clarisse Dumont, Guangping Li, Julien Castel, Serge Luquet, Giuseppe Gangarossa

**Author notes:** Correspondence to Giuseppe Gangarossa (, @PeppeGanga).

## Abstract

The regulation of food intake and energy balance relies on the dynamic integration of exteroceptive and interoceptive signals monitoring nutritional, metabolic, cognitive and emotional states. The paraventricular thalamus (PVT) is a central hub that, by integrating sensory, metabolic and emotional states, may contribute to the regulation of feeding and homeostatic/allostatic processes. However, the underlying PVT circuits remain still elusive. Here, we aimed at unraveling the role of catecholaminergic (CA) inputs to the PVT in scaling feeding and metabolic efficiency. First, using region-specific retrograde disruption of CA projections, we show that PVT CA inputs mainly arise from the hindbrain, notably the locus coeruleus (LC) and the nucleus tractus solitarius (NTS). Second, taking advantage of integrative calorimetric measurements of metabolic efficiency, we reveal that CA inputs to the PVT scale adaptive feeding and metabolic responses in environmental, behavioral, physiological and metabolic stress-like contexts. Third, we show that hindbrain^TH^→PVT inputs contribute in modulating the activity of PVT as well as lateral (LH) and dorsomedial (DMH) hypothalamic neurons.

In conclusion, this study, by assessing the key role of CA inputs to the PVT in scaling homeostatic/allostatic regulations of feeding patterns, reveals the integrative and converging hindbrain^TH^→PVT paths that contribute to whole-body metabolic adaptations in stress-like contexts.

**Key points:**

1. The paraventricular thalamus (PVT) is known to receive projections from the hindbrain. Here, we confirm and further extend current knowledge on the existence of hindbrain^TH^→PVT catecholaminergic (CA) inputs, notably from the locus coeruleus (LC) and the nucleus tractus solitarius (NTS), with the NTS representing the main source.
2. Disruption of hindbrain^TH^→PVT inputs contribute to the modulation of PVT-neurons activity.
3. Hindbrain^TH^→PVT inputs scale feeding strategies in environmental, behavioral, physiological and metabolic stress-like contexts.
4. Hindbrain^TH^→PVT inputs participate in regulating metabolic efficiency and nutrient partitioning in stress-like contexts.
5. Hindbrain^TH^→PVT, directly and/or indirectly, contribute in modulating the downstream activity of lateral (LH) and dorsomedial (DMH) hypothalamic neurons.

## Introduction

In mammals, the regulation of food intake intricately relies on the orchestration of several signals mirroring the dynamic integration of interoceptive and exteroceptive (environment) states (Sweeney & Yang, 2017). Indeed, emotional states (stress, anxiety, motivation), by modulating the activity of central neuronal hubs, thoroughly scale the regulation of food intake and the establishment of feeding habits (Ulrich-Lai *et al*., 2015). This is particularly evident in stress-related contexts where emotional states compete with the homeostatic regulation of feeding (Maniam & Morris, 2012; Herzog, 2020), thereby leading to feed-forward allostatic adaptations (*stability through changes*) which may culminate in psychiatric and metabolic disorders. These observations support the idea that emotional and homeostatic states may share similar, although not identical, neuronal networks (Sweeney & Yang, 2017). Nonetheless, the systems underlying the functional connection between these states remain poorly understood.

Emerging evidence strongly suggests that the paraventricular nucleus of the thalamus (PVT), a dorsal midline thalamic relay, may represent a functional *hot-spot* where interoceptive and exteroceptive stimuli may converge to orchestrate the selection of adaptive and appropriate behavioral responses aimed at regulating homeostatic and cognitive processes (Penzo & Gao, 2021). Given their elaborated connectivity (Kirouac, 2015), PVT excitatory (glutamate) neurons are well positioned to serve as functional integrators of orexigenic and anorexigenic stimuli (Ong *et al*., 2017; Meffre *et al*., 2019), glucose fluctuations (Labouèbe *et al*., 2016; Kessler *et al*., 2021), learning and memory processes (Do-Monte *et al*., 2015; Penzo *et al*., 2015) as well as emotional states (Heydendael *et al*., 2011; Barson *et al*., 2020; Pliota *et al*., 2020). This plethora of PVT-related brain functions highly relies on different afferent projections which, by carrying distinct neurochemical information, synergistically scale the activity of PVT-neurons and their downstream connected regions. Among the different afferents, the PVT also harbors a dense plexus of catecholaminergic (CA) fibers mainly arising from the hindbrain (Schroeter *et al*., 2000; Beas *et al*., 2018; Sofia Beas *et al*., 2020; Wang *et al*., 2021b) and only few scattered CA fibers from the hypothalamus (Li *et al*., 2014). In addition, recent reports have indicated that brain CA (norepinephrine, NE, and/or dopamine, DA), by modulating different homeostatic functions (*i.e.* wakefulness, arousal, glucoprivation-induced food seeking), may represent key neuromodulators of the PVT (Beas *et al*., 2018; Sofia Beas *et al*., 2020; Wang *et al*., 2021b). Moreover, brain CA are also important contributors to the regulation of stress-like responses (Valentino & Van Bockstaele, 2008; Kvetnansky *et al*., 2009) which ultimately impact, directly and/or indirectly, on feeding patterns and behaviors (Xu *et al*., 2019; Qu *et al*., 2020).

Indeed, the PVT has already been involved in food-seeking behaviors mostly associated to positive (motivation, reward, reinforcement) or negative valance (Labouèbe *et al*., 2016; Otis *et al*., 2017, 2019; Do-Monte *et al*., 2017; Wang *et al*., 2021a; Engelke *et al*., 2021) as well as in stress and emotional arousal (Hsu *et al*., 2014). However, whether and how the PVT and its afferent CA inputs may contribute to the regulation of food intake and metabolic efficiency in stress-related contexts remain to be fully established.

In order to dissect the contribution of PVT CA inputs in the regulation of food intake, we suppressed local CA inputs by microinjecting the neurotoxin 6-OHDA into the PVT and performed several experiments aimed at revealing the adaptive strategies associated to the regulation of feeding and energy balance. Here, we demonstrate that CA inputs to the PVT exerted a modulatory role on food intake and metabolic efficiency specifically in stress-related contexts since no major alterations were detected in basal conditions. Notably, we found that 6-OHDA^PVT^-lesioned mice, following both exteroceptive (environmental) and interoceptive (body energy) stressors, showed enhanced food intake and metabolic efficiency.

Altogether, our results reveal a novel neuronal network by which stressors impinge on the regulatory allostatic processes underlying feeding behaviors, metabolic efficiency and nutrient partitioning to scale stress-associated adaptive responses.

## Materials and methods

### Ethics statement and animals

All experiments were approved by the Animal Care Committee of the Université Paris Cité (APAFiS #35585 and #11003) and carried out following the 2010/63/EU directive. 8-14 weeks old male C57Bl/6J mice (Janvier, Le Genest St Isle, France) were used and housed in a room maintained at 22 +/-1 °C, with a light period from 7h00 to 19h00. Regular chow diet (3.24 kcal/g, reference SAFE^®^ A04, Augy, France) and water were provided *ad libitum* unless otherwise stated.

### Stereotaxic microinjections for viral tracing studies and 6-OHDA-induced catecholaminergic denervation

Mice were anaesthetized with isoflurane (3.5% for induction, 1.5% for maintenance), administered with Buprécare^®^ (buprenorphine 0.3 mg) and Ketofen^®^ (ketoprofen 100 mg), and placed on a stereotactic frame (Model 940, David Kopf Instruments). During surgery, body temperature was maintained at 37°C.

6-OHDA-HCl (Sigma-Aldrich, #H4381) was dissolved in a saline solution (NaCl 0.9% w/v) containing 0.02% of ascorbic acid at a final concentration of 6 µg/µl. Animals were randomly assigned to either 6-OHDA or vehicle microinjections. A volume of 0.35 µl of 6-OHDA or vehicle (0.02% ascorbic acid) was injected into the PVT (L= 0.0; AP= -1.46; V= -2.8, mm) at an infusion rate of 0.05 µl/min. The injection needle was carefully removed after waiting 5 minutes at the injection site. 24-hrs after, animals were re-administered with Buprécare^®^ and Ketofen^®^, and had facilitated access to jelly food (DietGel Boost #72-04-5022, Clear H_2_O) for 2 consecutive days. Recovery from surgery was monitored during 3-5 days post-surgery. Animals were allowed to recover for 3-4 weeks before any experimental evaluation.

pAAV-CAG-tdTomato (titer ≥ 1×10¹³ vg/mL) was a gift from Edward Boyden (Addgene viral prep #59462-AAV9; http://n2t.net/addgene:59462; RRID:Addgene_59462). A volume of 0.20 µl of pAAV-CAG-tdTomato was injected into the PVT (L= 0.0; AP= -1.46; V= -2.8, mm) at an infusion rate of 0.05 µl/min. The injection needle was carefully removed after waiting 5 minutes at the injection site. Viral expression was evaluated 4 weeks after microinjection.

### Indirect calorimetry and metabolic efficiency analysis

Indirect calorimetry was performed as previously described (Berland *et al*., 2022). Mice were monitored for whole energy expenditure (EE), O_2_ consumption and CO_2_ production, respiratory exchange rate (RER=VCO_2_/VO_2_), fatty acid oxidation (FAO), and locomotor activity using calorimetric cages with bedding, food and water (Labmaster, TSE Systems GmbH, Bad Homburg, Germany). Ratio of gases was determined through an indirect open circuit calorimeter (Arch *et al*., 2006; Even & Nadkarni, 2012). This system monitors O_2_ and CO_2_ concentration by volume at the inlet ports of a tide cage through which a known flow of air is being ventilated (0.4 L/min) and compared regularly to a reference empty cage. For optimal analysis, the flow rate was adjusted according to the animal body weights to set the differential in the composition of the expired gases between 0.4-0.9% (Labmaster, TSE Systems GmbH, Bad Homburg, Germany). The flow was previously calibrated with O_2_ and CO_2_ mixture of known concentrations (Air Liquide, S.A. France). O_2_ consumption and CO_2_ production were recorded every 15 min for each animal during the entire experiment. Whole energy expenditure (EE) was calculated using the Weir equation for respiratory gas exchange measurements. Food consumption was measured as the instrument combines a set of highly sensitive feeding sensors for automated online measurements. Mice had access to food and water *ad libitum*. To allow measurement of every ambulatory movement, each cage was embedded in a frame with an infrared light beam-based activity monitoring system with online measurement at 100 Hz. The sensors for gases and detection of movements operated efficiently in both light and dark phases, allowing continuous recording. When required, the inversion of circadian light/dark cycles was programmed using the Labmaster software.

Body mass composition was analyzed using an Echo Medical systems’ EchoMRI (Whole Body Composition Analyzers, EchoMRI, Houston, USA), according to manufacturer’s instructions. Readings of body composition were given within 1 min. Data analysis was performed on Excel XP using extracted raw values of VO_2_ consumed (expressed in ml/h), VCO_2_ production (expressed in ml/h), and energy expenditure (kcal/h).

### Novelty-suppressed feeding (NSF)

After an overnight fasting, mice were placed in a cage (40×40×40 cm) with a single regular chow pellet in the center. The latency (time in seconds) to eat was scored. To measure food intake, food consumption was evaluated 60 minutes after the beginning of the test.

### Open field (OF)

Spontaneous exploratory behavior was monitored in an open field (40×40×40 cm, BIOSEB) for 20 min, video-tracked and analyzed using the SMART software (BIOSEB). The open field was wiped with 70% ethanol between sessions.

### Infrared temperature measurements

Thermogenesis was visualized using a high-resolution infrared camera (FLIR E8; FLIR Systems, Portland, OR, USA). To measure the temperature (°C) of the brown adipose tissue (BAT, interscapular regions), lower back and tail, images were captured before and after the open field (OF) test. Infrared thermography images were analyzed using the FLIR TOOLS software.

### Food intake following food or water deprivation

In two distinct experiments we measured food intake following either an overnight fasting (food deprivation) or water deprivation.

#### Overnight fasting

Mice were first weighted in the morning following an overnight fasting to ensure the loss of body weight. Then, they were exposed to pre-weighted chow pellets. Food intake was measured at the following time points: 30 min, 1h, 2h, 3h, 4h.

#### Overnight water deprivation

Mice were first weighted in the morning following an overnight water deprivation to ensure the loss of body weight. Then, they were exposed to a calibrated drinking bottle and pre-weighted chow pellets. Water consumption and food intake were measured at the following time points: 30 min, 1h, 2h, 3h, 4h.

### Food intake induced by 2-DG and ghrelin

Mice were handled and injected with vehicle during 3 consecutive days before drugs administration. After this habituation period, they were administered with ghrelin (#1465, Tocris, 0.5 mg/kg, i.p.) or 2-DG (#14325, Cayman Chemical, 500 mg/kg, i.p.) and exposed to chow pellets 30 min after the injections. Food intake was measured during 3 hours. For 2-DG treated mice, blood glucose (counterregulatory response) was measured from the vein tail using a glucometer (Menarini Diagnotics, Rungis, France) at 0 and 30 min.

### Glucose dynamics

#### Oral glucose tolerance test (OGTT)

Animals were fasted 5 hours before receiving a bolus of glucose solution (2 g/kg) by gavage. Blood glucose (hyperglycemia) was measured from the vein tail using a glucometer (Menarini Diagnotics, Rungis, France) at 0, 5, 10, 15, 30, 60, 90, and 120 min.

#### Insulin tolerance test (ITT)

Animals were fasted 5 hours before receiving an injection of insulin (0.5 U/kg, Novo Nordisk, i.p.). Blood glucose (hypoglycemia) was measured from the vein tail at 0, 5, 10, 15, 20, 30, 60, 90, and 120 min.

### Tissue preparation and immunofluorescence

Animals were injected with pentobarbital (500 mg/kg, i.p., Sanofi-Aventis, France). Once anaesthetized, they were transcardially perfused with cold (4°C) PFA 4% for 5 minutes. Brains were collected, put overnight in PFA 4% and then stored in PBS (4°C). 40 µm-thick sections were sliced with a vibratome (Leica VT1000S, France) and stored in a cryoprotective solution at -20 °C until immunofluorescence investigations. Immunofluorescence on brain slices was performed as previously described (Gangarossa *et al*., 2019; Berland *et al*., 2020).

Briefly, sections were processed as it follows. Day 1: free-floating sections were rinsed in Tris-buffered saline (TBS; 0.25 M Tris and 0.5 M NaCl, pH 7.5), incubated for 5 min in TBS containing 3% H_2_O_2_ and 10% methanol, and then rinsed three times for 10 min each in TBS. After 15 min incubation in 0.2% Triton X-100 in TBS, sections were rinsed three times in TBS again. Slices were then incubated 48 hrs at 4°C with the following primary antibodies: rabbit anti-cFos (1:1000, Synaptic Systems, #226 003), rabbit anti-TH (1:1000, Millipore, #AB153) or rat anti-DAT (1:500, Millipore, #MAB369). Sections were rinsed three times for 10 min in TBS and incubated for 60 min with secondary donkey anti-rabbit Cy3 AffiniPure (1:1000, Jackson ImmunoResearch, #711-165-152) or donkey anti-rat Cy3 AffiniPure (1:1000, Jackson ImmunoResearch, #712-165-153). Sections were rinsed for 10 min twice in TBS, stained with DAPI (10 min) and rinsed in TB (0.25 M Tris) before mounting.

Acquisitions were performed with a confocal microscope (Zeiss LSM 510). The objective (10X) and the pinhole setting remained unchanged during the acquisition of a series for all images. Depending on the extension of the region of interest, either single or mosaic acquisitions were conducted. Quantification of immunoreactive cells (cFos- or TH-positive neurons) was performed using the cell counter plugin of the ImageJ software taking as standard reference a fixed threshold of fluorescence. For cell counting, three (TH) or two (cFos) rostro-caudal levels for each brain region were used. Quantifications of immunopositive neurons were averaged between hemispheres and then summed for consecutive brain slices.

### Statistics

All data are presented as mean ± SD. Statistical tests were performed with Prism 7 (GraphPad Software, La Jolla, CA, USA). Detailed statistical analyses are listed in the **Statistical Summary Table**. Normality was assessed by the D’Agostino-Pearson test. Depending on the experimental design, data were analyzed using either Student t-test (paired or unpaired) with equal variances, One-way ANOVA or Two-way ANOVA. The significance threshold was automatically set at p<0.05. ANOVA analyses were followed by Bonferroni *post hoc* test for specific comparisons only when overall ANOVA revealed a significant difference (at least p<0.05).

## Results

### Catecholaminergic inputs modulate the activity of PVT-neurons

To study whether catecholaminergic (TH-positive) inputs to the PVT participate/contribute to the regulation of food intake, we decided to ablate catecholamine (CA) fibers projecting to the PVT by locally microinjecting 6-OHDA (**Figure 1A**), a neurotoxin reuptaken by DAT- and/or NET-expressing terminals. Since the PVT extends throughout the midline thalamic axis, we decided to mainly focus on the mid-posterior PVT as this region has been shown to be involved in stress and stress-induced hypophagia (Heydendael *et al*., 2011; Barson *et al*., 2020; Barrett *et al*., 2021). Indeed, 6-OHDA was able to strongly reduce TH immunostaining in the PVT (**Figure 1B**), with few remaining terminals most likely corresponding to NET-negative TH fibers (terminals releasing adrenaline and/or devoid of monoamine transporters) since the PVT does not seem to contain DAT-positive terminals as compared to DAT-rich regions such as the tail of the striatum [TS, (Gangarossa *et al*., 2013; Valjent & Gangarossa, 2021)] and the central amygdala (**Suppl. Figure 1A**, https://doi.org/10.6084/m9.figshare.19683228.v1).

**Figure 1:**
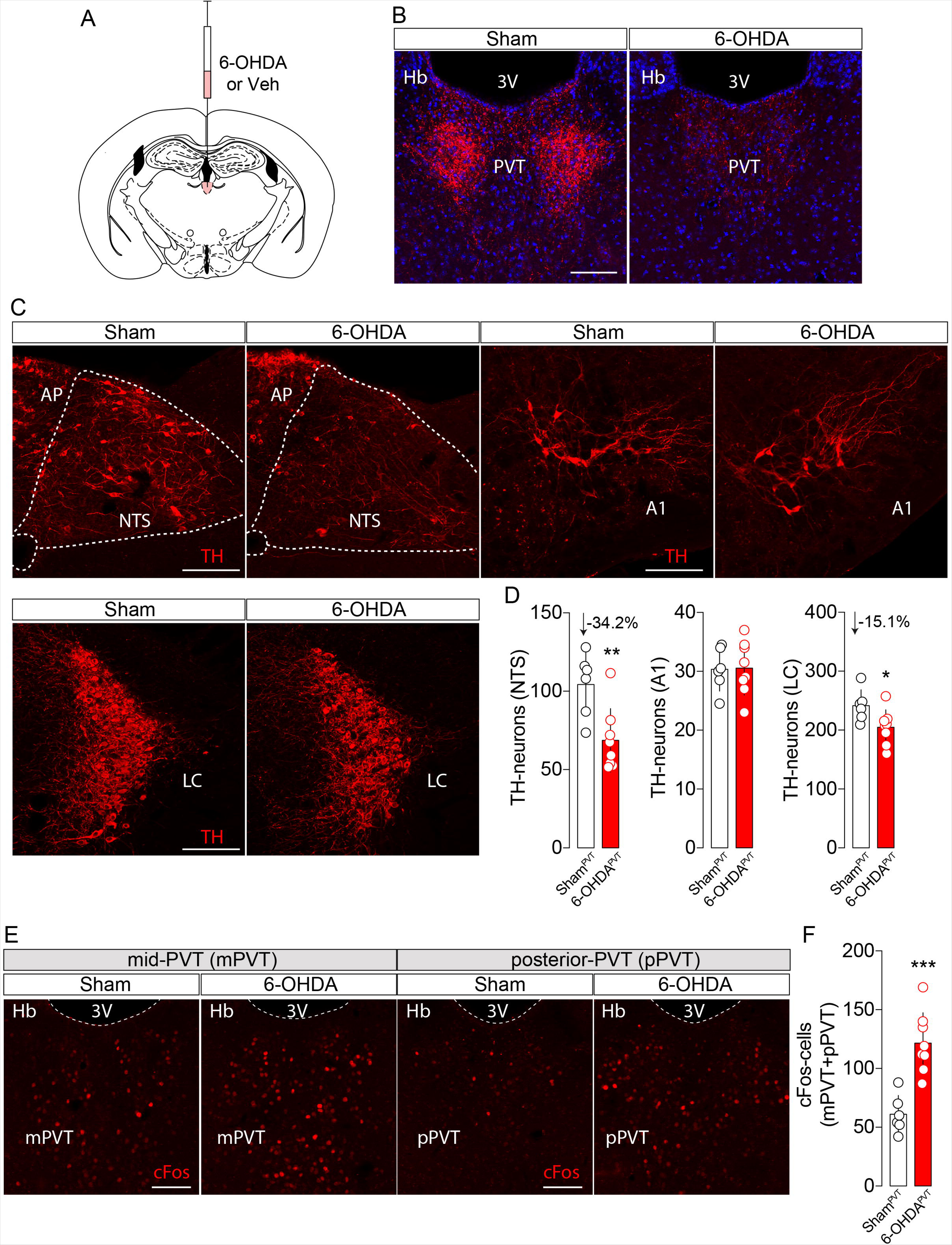
Retrograde ablation of hindbrain catecholaminergic inputs contribute to PVT activity. (**A**) Drawing represents the microinjection of 6-OHDA or Vehicle (Veh) in the mid-posterior PVT. (**B**) Immunofluorescence detection of tyrosine hydroxylase (TH) within the PVT in Sham^PVT^ and 6-OHDA^PVT^ mice. Scale bar: 150 µm. (**C**) Immunofluorescence detection of TH in PVT-projecting hindbrain regions [nucleus tractus solitarius (NTS), area A1 and locus coeruleus (LC)] in Sham^PVT^ and 6-OHDA^PVT^ mice. Scale bars: 150 µm. (**D**) Quantification of TH-positive neurons in the NTS, A1 and LC. Statistics: * p<0.05, ** p<0.01, 6-OHDA^PVT^ *vs* Sham^PVT^ mice. (**E**) Immunofluorescence detection of cFos-positive neurons in the mid-posterior PVT (mPVT and pPVT). Scale bars: 150 µm. (**F**) Quantification of cFos-positive cells in the mid-posterior PVT. Statistics: *** p<0.001, 6-OHDA^PVT^ *vs* Sham^PVT^ mice. Data are presented as mean ± SD. For statistical details see **Statistical Summary Table**.

We therefore examined the regional sources of our 6-OHDA-induced degeneration of TH-afferents. We focused on putative noradrenergic TH-positive projecting neurons since the PVT do not receive dopaminergic inputs from the midbrain (SNc and VTA) (Li *et al*., 2014) and it harbors only minor, if any, scattered DAT-fibers (García-Cabezas *et al*., 2009; Clark *et al*., 2017). Immunofluorescence analysis revealed a significant reduction of TH-neurons in the nucleus tractus solitarius (NTS) and the locus coeruleus (LC), in line with the presence of *Slc6a2* (NET)-cathecolaminergic neurons in these nuclei (Schroeter *et al*., 2000; Zhang *et al*., 2021) and the sensitivity of these neurons to 6-OHDA (Szot *et al*., 2012b; Lin *et al*., 2013). However, we observed a more robust reduction of PVT-projecting TH-neurons in the NTS (−34.2%) compared to the LC (−15.1%) (**Figure 1C, D**). No differences were observed in the A1 area of the hindbrain (**Figure 1C, D**) as well as in the hypothalamus (**Suppl. Figure 1B**, https://doi.org/10.6084/m9.figshare.19683228.v1).

Then we investigated whether the reduction in local TH-afferents was followed by the modulation of PVT-neurons activity. Since the PVT shows higher activity during wakefulness (Ren *et al*., 2018), mice were perfused 1h before the onset of the dark phase (spontaneous feeding period) and at basal conditions. Using cFos as a molecular proxy of cellular activity, we observed a significant increase in cFos-positive cells in the PVT of 6-OHDA^PVT^ compared to Sham^PVT^ mice (**Figure 1E, F**). This set of results suggests that a subset of hindbrain CA inputs (hindbrain^TH^→PVT projections) may serve as modulators of PVT-neurons activity.

### Catecholaminergic inputs to the PVT contribute to novelty-induced hypophagia

Following 3-4 weeks from 6-OHDA microinjections, no major differences in body weight were detected in 6-OHDA^PVT^ compared to Sham^PVT^ mice (group-housed animals, **Figure 2A**). In order to study the role of PVT catecholaminergic inputs in the regulation of feeding patterns, 6-OHDA^PVT^ and Sham^PVT^ mice were single-housed (**Figure 2B**). Environmental and social isolation, as triggered by single housing, represent behavioral/environmental stress-like allostatic stimuli (Lee *et al*., 2020, 2021) which can trigger a transient reduction of food intake (Takatsu-Coleman *et al*., 2013; Benfato *et al*., 2022). Interestingly, we observed that 6-OHDA^PVT^ mice were less sensitive to single housing-induced hypophagia compared to Sham^PVT^ mice (**Figure 2C**). This phenotype prompted us to investigate whether catecholaminergic inputs to the PVT were important in driving feeding patterns and metabolic adaptations to exposure to other behavioral/environmental challenges.

**Figure 2:**
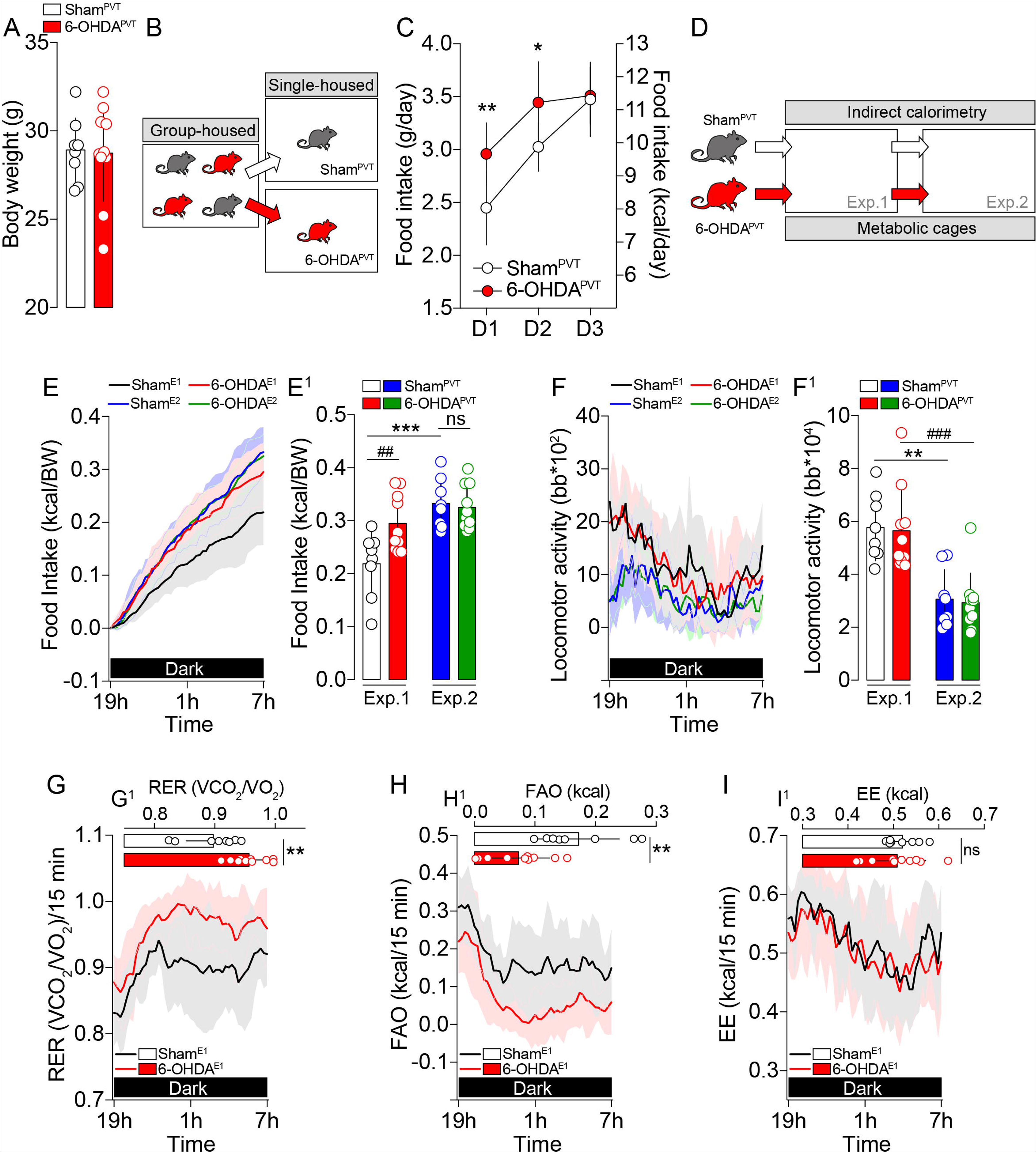
Hindbrain^TH^→PVT inputs participate to novelty-induced hypophagia. (**A**) Body weight of Sham^PVT^ and 6-OHDA^PVT^ mice following 3-4 weeks from 6-OHDA or Veh microinjection in the PVT. (**B**) Experimental design indicating the transition from grouped to singled housing. (**C**) Food intake (g/day and kcal/day) during the first three days of isolation (D1 to D3). Statistics: * p<0.05, ** p<0.01, 6-OHDA^PVT^ *vs* Sham^PVT^ mice. (**D**) Investigation of food intake and metabolic efficiency using metabolic cages during two consecutive exposures to a novel environment. (**E**) Food intake during the dark phase (spontaneous eating) in Sham^PVT^ and 6-OHDA^PVT^ mice during two consecutive exposures to a novel environment. (**E^1^**) Cumulative food intake. Statistics: *** p<0.01, Sham^E2^ *vs* Sham^E1^, ^##^ p<0.01, 6-OHDA^E1^ *vs* Sham^E1^ groups. (**F**) Spontaneous locomotor activity (beam breaks, bb) during the dark phase in Sham^PVT^ and 6-OHDA^PVT^ mice during two consecutive exposures to a novel environment. (**F^1^**) Cumulative locomotor activity. Statistics: ** p<0.01, Sham^E2^ *vs* Sham^E1^; ^###^ p<0.001, 6-OHDA^E2^ *vs* 6-OHDA^E1^ groups. Note that both experimental groups showed similar degrees of habituation (reduced locomotor activity during the dark period). Measurements of the respiratory exchange ratio (RER, **G**), fatty acid oxidation (FAO, **H**), and energy expenditure (EE, **I**) in Sham^PVT^ and 6-OHDA^PVT^ mice during the first exposure (Exp.1) to a novel environment. (**G^1^-1^1^**) Averaged RER, FAO and EE during the dark phase. Statistics: ** p<0.01, 6-OHDA^E1^ *vs* Sham^E1^. Data are presented as mean ± SD. For statistical details see **Statistical Summary Table**.

After 2 weeks of habituation to single-housing, we analyzed the metabolic efficiency of 6-OHDA^PVT^ and Sham^PVT^ mice exposed to a novel environment during two consecutive exposures during the dark period (spontaneous feeding period, **Figure 2D**). While Sham^PVT^ mice transiently (Exp.1) showed a reduction in food intake during the dark period (**Figure 2E, E^1^**), 6-OHDA^PVT^ mice were again less sensitive to novelty-induced hypophagia. No differences in food intake were measured during Exp.2 period between the two groups, indicating a rapid restoration of homeostatic regulations associated to environmental habitation (**Figure 2E, E^1^**). This phenotype was not associated to changes in locomotor activity as indicated by the similar patterns of exploration (Exp.1, novelty) and habituation (Exp.2) (**Figure 2F, F^1^**). We also measured key whole-body metabolic parameters such as the respiratory exchange ratio (RER, indicative of the energy substrates used, RER=∼1 for carbohydrates, RER=∼0.7 for lipids), fatty acid oxidation (FAO) and energy expenditure (EE) during the exposure to the novel environment (Exp.1). Compared to Sham^PVT^ mice, 6-OHDA^PVT^ animals showed increased RER (**Figure 2G, G^1^**) and decreased FAO (**Figure 2H, H^1^**), mirroring the changes in food intake and indicating a shift of energy substrates (carbohydrates and lipids) use during exposure to a novel environment. However, these adaptations did not impact on energy expenditure (**Figure 2I, I^1^**), suggesting that nutrient partitioning (Joly-Amado *et al*., 2012), rather than total energy balance, was affected by the loss of hindbrain^TH^→PVT fibers.

One may wonder whether the absence of novelty-induced hypophagia in 6-OHDA^PVT^ mice may be associated to enhanced perception and/or reward value of food. To investigate this aspect, Sham^PVT^ and 6-OHDA^PVT^ mice were intermittently (1h/day) exposed to high-fat high-sugar (HFHS) diet during two consecutive days. No differences in HFHS food intake were observed between groups (**Suppl. Figure 2A**, https://doi.org/10.6084/m9.figshare.19683228.v1), suggesting intact food palatability, perception and preference. These results suggest that hindbrain^TH^→PVT fibers represent an important node for the integration of homeostatic regulations.

Exposure to novel environments can lead to the occurrence of anxiogenic traits and novelty-triggered thermogenic adaptations (Lecorps *et al*., 2016) which may impact on, and therefore confound, the mechanisms underlying feeding strategies. Thus, we performed an open field test (OF, **Figure 3A**) to evaluate both anxiety and thermogenic adaptations. We observed no significant differences in anxiety-like parameters between Sham^PVT^ and 6-OHDA^PVT^ animals (**Figure 3B-F**). Moreover, both groups showed similar thermogenic enhancements in the brown adipose tissue (BAT), the lower back and the tail (**Figure 3G-J**). These results indicate that PVT-projecting TH-afferents modulate feeding patterns and metabolic efficiency independently from affective (anxiety) and thermogenic adaptations.

**Figure 3:**
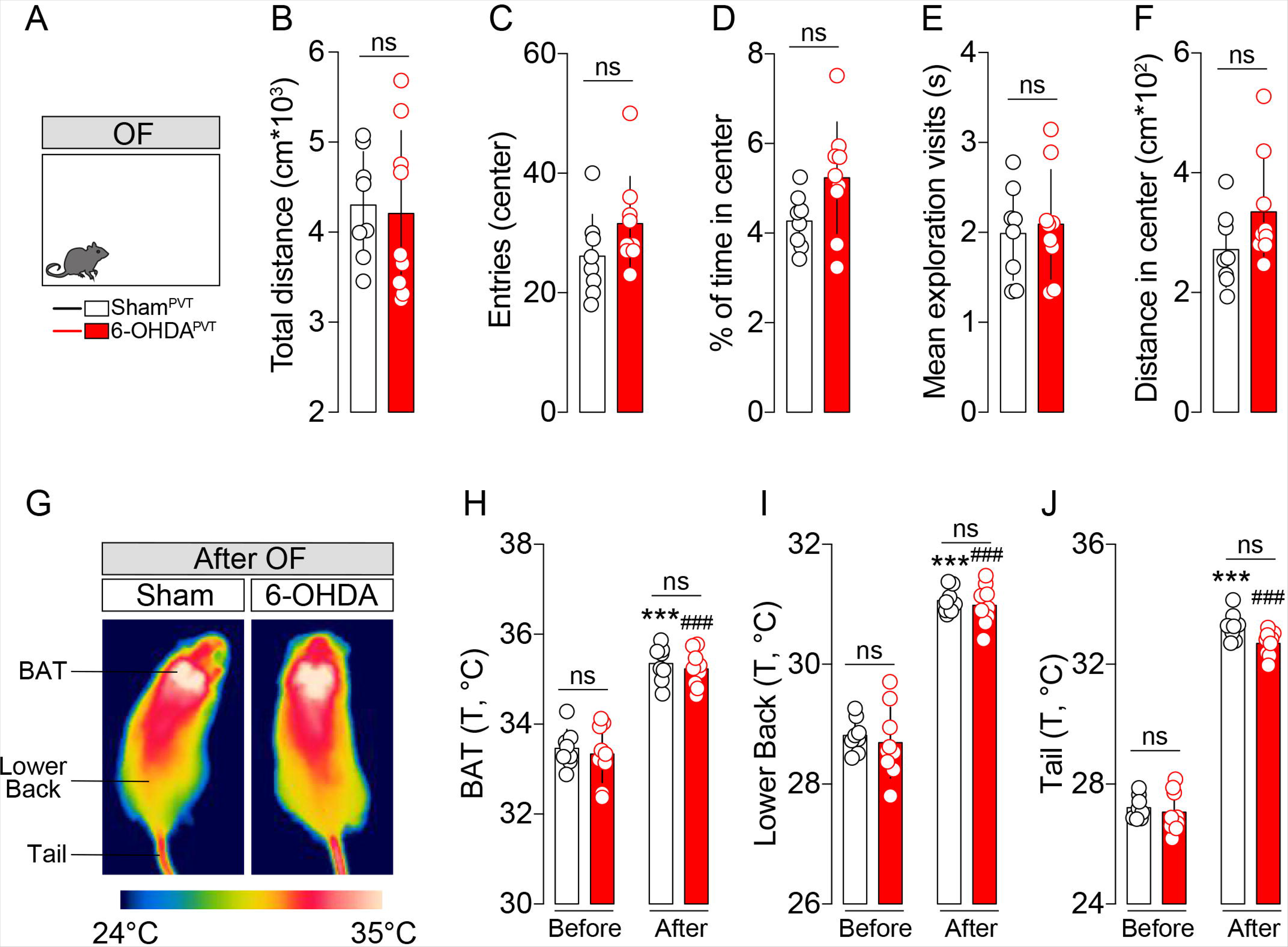
Hindbrain^TH^→PVT inputs do not alter novelty-induced anxiety and thermogenesis. (**A**) Drawing represents the open field (OF) test. (**B-F**) Parameters measured during a 20 min OF test: total distance, number of entries in the center, % of time in the center, mean exploration visits to the center, distance in the center. (**G**) Infrared thermographic images of animals after the OF test (20 min). (**H-J**) Temperature (°C) of the brown adipose tissue (BAT), lower back and tail in Sham^PVT^ and 6-OHDA^PVT^ mice before and after the OF test. Statistics: *** p<0.001, After *vs* Before OF test (Sham^PVT^ mice); ^###^ p<0.001, After *vs* Before OF test (6-OHDA^PVT^ mice). No differences were detected between Sham^PVT^ and 6-OHDA^PVT^ mice. Data are presented as mean ± SD. For statistical details see **Statistical Summary Table**.

Feeding patterns and metabolic efficiency highly depend on circadian rhythms and strong functional interactions exist between circadian rhythms, feeding and energy balance (Challet, 2019). Moreover, the PVT, which is bidirectionally connected with the suprachiasmatic nucleus (SCN) (Peng & Bentivoglio, 2004; Colavito *et al*., 2015), has been pointed as a contributor to circadian cycles and its activity varies depending on active/inactive phases (Colavito *et al*., 2015; Kirouac, 2015). Thus, we decided to investigate whether PVT catecholaminergic inputs participated to the adaptive metabolic strategies occurring during circadian challenges by inverting the light/dark cycle (**Figure 4A**). First, before inverting the light/dark cycle, we confirmed (**Figure 2**) that exposure to the new environment was associated to reduced novelty-induced hypophagia in 6-OHDA^PVT^ compared to Sham^PVT^ mice (**Suppl. Figure 3A, B**, https://doi.org/10.6084/m9.figshare.19683228.v1). Then, after habituation, light/dark cycles were inverted. When measuring food intake, we observed that both experimental groups rapidly shifted and adapted toward the new light/dark schedule (average of first 2 days of standard and inverted cycles) (**Figure 4B, B^1^**). In fact, despite the adaptive shift (**Figure 4B, B^1^**), no differences were observed in the cumulative food intake (**Figure 4C**). Moreover, we also detected a similar pattern of circadian adaptation in the EE profile of both groups (**Figure 4D, D^1^**), with an increased EE in the 7h-19h inverted period (iDark) and a decreased EE in the 19h-7h inverted period (iLight) (**Figure 4E, E^1^**). Interestingly, we found a significant difference in FAO and RER. In particular, while Sham^PVT^ animals rapidly adapted to the inverted light/dark cycle (**Figure 4F**), 6-OHDA^PVT^ mice did not show significant changes in FAO during the 7h-19h inverted period (iDark) (**Figure 4F^1^, G**), whereas both groups showed increased FAO during the 19h-7h inverted period (iLight) (**Figure 4G^1^**). In line with this shift in energy substrates partitioning, measurements of RER indicated an impaired adjustment of metabolic efficiency during the iDark period (7h-19h) in 6-OHDA^PVT^ mice (**Figure 4H, H^1^, I, I^1^**).

**Figure 4:**
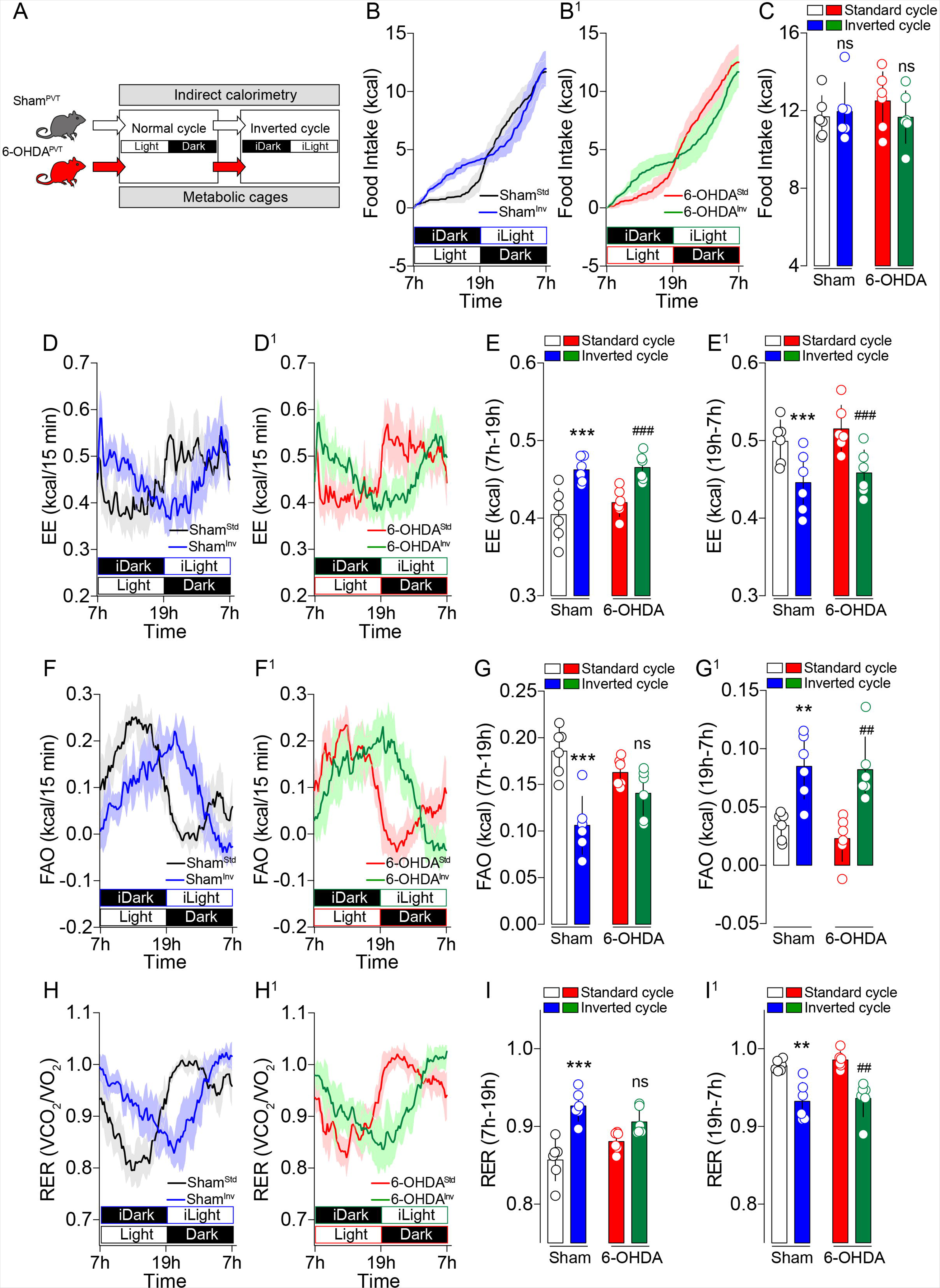
Hindbrain^TH^→PVT inputs contribute to the circadian adaptation of metabolic efficiency. (**A**) Drawing indicates the experimental procedure used to study feeding and metabolic adaptations during an inverted cycle (transition from Light-to-Dark to Dark-to-Light). (**B, B^1^**) Food intake in Sham^PVT^ (**B**) and 6-OHDA^PVT^ (**B^1^**) mice during the standard and inverted cycles. Note how the temporal dynamics of feeding change during the inverted cycle. (**C**) Cumulative food intake in Sham^PVT^ and 6-OHDA^PVT^ mice during the standard and inverted cycles. (**D, D^1^**) Temporal dynamics of EE adaptations in Sham^PVT^ (**D**) and 6-OHDA^PVT^ (**D^1^**) mice during the standard and inverted cycles. (**E, E^1^**) Averaged EE in Sham^PVT^ and 6-OHDA^PVT^ mice according to matched inverted phases (7h-19h, **E**, and 19h-7h, **E^1^**). Statistics: *** p<0.001, Inverted *vs* Standard cycle (Sham^PVT^ mice); ^###^ p<0.001, Inverted *vs* Standard cycle (6-OHDA^PVT^ mice). (**F, F^1^**) Temporal dynamics of FAO adaptations in Sham^PVT^ (**F**) and 6-OHDA^PVT^ (**F^1^**) mice during the standard and inverted cycles. (**G, G^1^**) Averaged FAO in Sham^PVT^ and 6-OHDA^PVT^ mice according to matched inverted phases (7h-19h, **G**, and 19h-7h, **G^1^**). Statistics: *** p<0.001, ** p<0.01, Inverted *vs* Standard cycle (Sham^PVT^ mice); ^##^ p<0.01, Inverted *vs* Standard cycle (6-OHDA^PVT^ mice). (**H, H^1^**) Temporal dynamics of RER adaptations in Sham^PVT^ (**H**) and 6-OHDA^PVT^ (**H^1^**) mice during the standard and inverted cycles. (**I, I^1^**) Averaged RER in Sham^PVT^ and 6-OHDA^PVT^ mice according to matched inverted phases (7h-19h, **I**, and 19h-7h, **I^1^**). Statistics: *** p<0.001, ** p<0.01, Inverted *vs* Standard cycle (Sham^PVT^ mice); ^##^ p<0.01, Inverted *vs* Standard cycle (6-OHDA^PVT^ mice). Data are presented as mean ± SD. For statistical details see **Statistical Summary Table**.

These results suggest that hindbrain^TH^→PVT inputs, although not involved in the adaptation of food intake to circadian challenges, are important in adjusting peripheral energy substrates utilization (*i.e* lipids and carbohydrates as revealed by RER and FAO) and overall metabolic efficiency.

### Catecholaminergic inputs to the PVT contribute to feeding under physiological and metabolic challenges

Next, we wondered whether hindbrain^TH^→PVT inputs guided feeding following physiological and metabolic stressors. First, to mimic a conflict between hunger and environmental stress, we decided to study food intake in overnight fasted mice in a novelty-induced hypophagia test. After an overnight fasting, both grouped showed a similar reduction in body weight and plasma glucose levels (**Suppl. Figure 4A, B**, https://doi.org/10.6084/m9.figshare.19683228.v1). Then mice underwent the novelty-suppressed feeding (NSF) test (**Figure 5A**). Sham^PVT^ and 6-OHDA^PVT^ mice showed similar latency to eat (**Figure 5B**), with a significant difference in food intake which was higher in 6-OHDA^PVT^ mice (**Figure 5C, D**). The NSF test (latency to eat) is mainly used to assess anxiety- and depressive-like phenotypes. Therefore, our results suggest that the increased food intake observed following PVT catecholaminergic ablation may not be confounded by potential alterations triggered by anxiety. This is in line with our above-mentioned observations using the open field test (**Figure 3A-F**).

**Figure 5:**
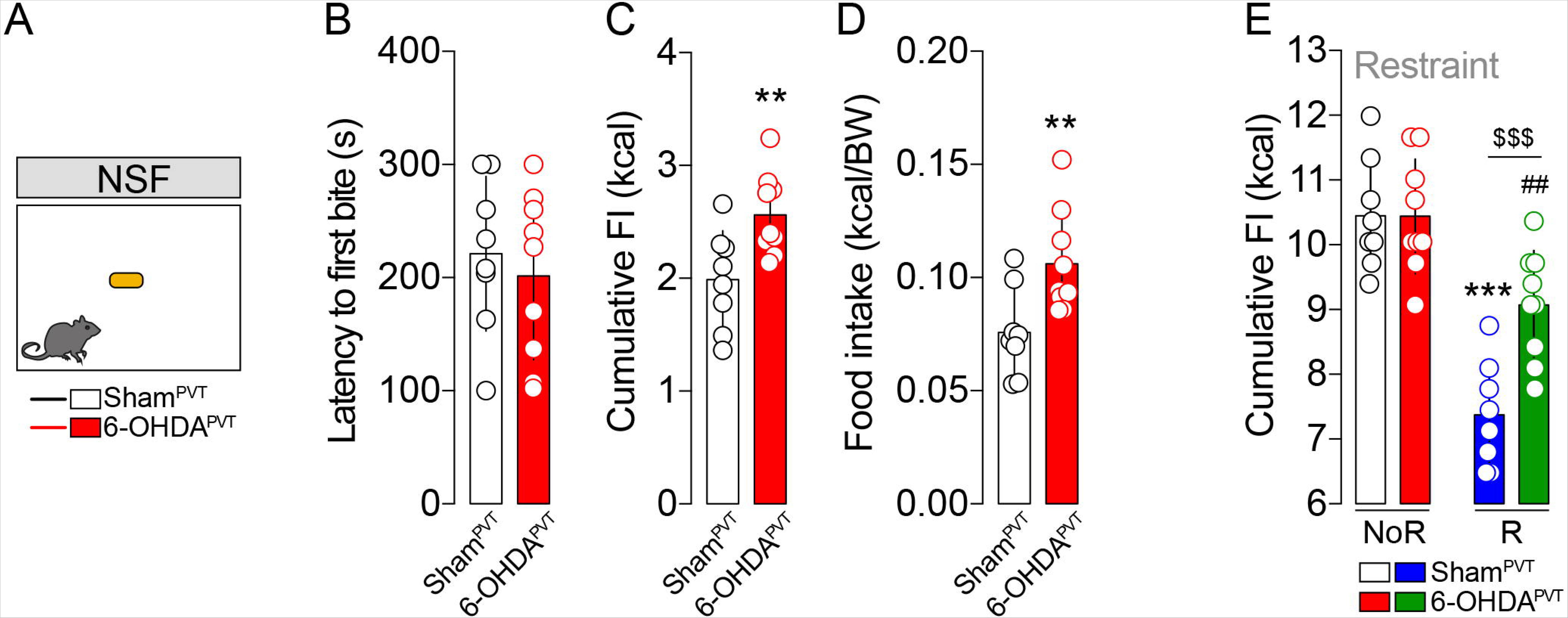
Hindbrain^TH^→PVT inputs scale feeding following physiological stressors. (**A**) Drawing represents the novelty-suppressed feeding (NSF) test in overnight fasted Sham^PVT^ and 6-OHDA^PVT^ mice. (**B**) Latency to eat (first bite) during the NSF test. (**C, D**) Food intake, total kcal (**C**) and normalized kcal/BW (**D**) during the NSF test. Statistics: ** p<0.01, 6-OHDA^PVT^ *vs* Sham^PVT^ mice. (**E**) Food intake in Sham^PVT^ and 6-OHDA^PVT^ mice after a restraint stress (immobilization). NoR: No Restraint (control conditions). R: Restraint. Statistics: *** p<0.001, Restraint *vs* No Restraint (Sham^PVT^); ^##^ p<0.01, Restraint *vs* No Restraint (6-OHDA^PVT^); ^$$$^ p<0.001, 6-OHDA^PVT^ *vs* Sham^PVT^ mice (Restraint). Data are presented as mean ± SD. For statistical details see **Statistical Summary Table**.

Then, we used an acute restraint (immobilization) paradigm which is known to alter metabolism and food intake (Rybkin *et al*., 1997; Vallès *et al*., 2000; Rabasa & Dickson, 2016). Sham^PVT^ and 6-OHDA^PVT^ mice underwent a 30 min acute restraint and food intake was measured during the dark period. Although both groups showed a significant reduction in food intake (**Figure 5E**), stress-induced hypophagia was significantly more pronounced in Sham^PVT^ compared to 6-OHDA^PVT^ mice (**Figure 5E**).

To further investigate the role of hindbrain^TH^→PVT inputs in scaling feeding, we used metabolic stressors to modulate food intake. First, Sham^PVT^ and 6-OHDA^PVT^ mice were administered with 2-deoxy-d-glucose (2-DG) which, in virtue of its glucoprivic effects (neuroglucopenia), elicits food consumption as well as the typical glucose counterregulatory response (CRR) (Pénicaud *et al*., 1986; Lewis *et al*., 2006). Moreover, PVT-neurons are highly sensitive to glucoprivation (Labouèbe *et al*., 2016). In this conditions, 6-OHDA^PVT^ mice consumed more chow food than Sham^PVT^ mice (**Figure 6A**), even though the magnitude of the glucose excursion as counterregulatory response was similar between groups (**Figure 6B**). This result suggests that hindbrain^TH^→PVT projections are required to fully express feeding adaptions to glucoprivic conditions but are dispensable for the autonomic control of glycogen breakdown and glucose production in CRR. In the same line, glucose clearance dynamics during an oral glucose tolerance test (OGTT, **Figure 6C**) or an insulin tolerance test (ITT, **Figure 6D**) were similar between sham and 6-OHDA^PVT^ mice, indicating that glucose metabolism and insulin sensitivity remained unaltered following the loss of hindbrain^TH^→PVT inputs.

**Figure 6:**
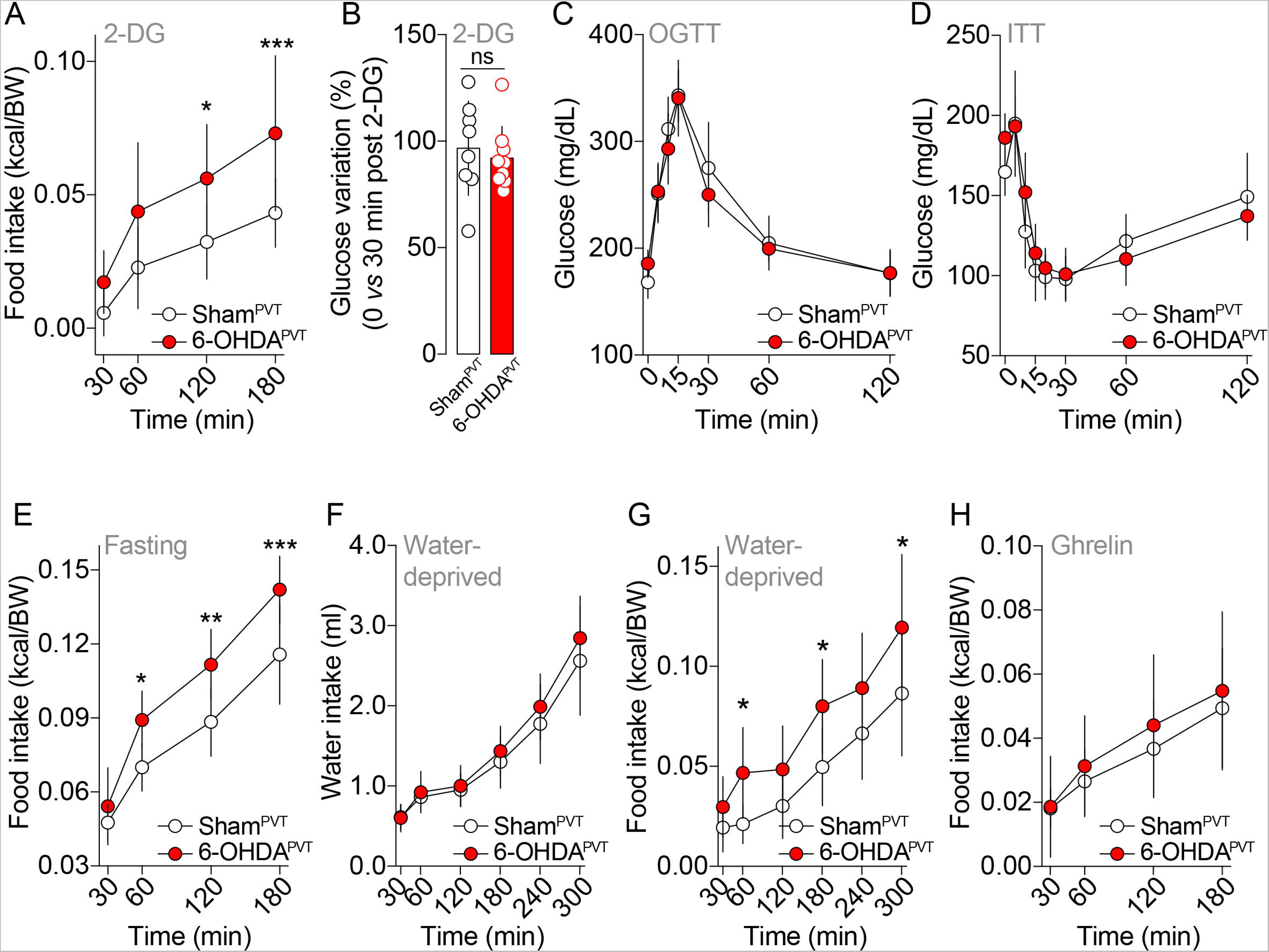
Hindbrain^TH^→PVT inputs scale feeding following metabolic stressors. (**A**) Food intake in Sham^PVT^ and 6-OHDA^PVT^ mice administered with 2-DG (500 mg/kg, i.p., neurogluopenia-induced feeding). (**B**) Glucose variation (%) following administration of 2-DG (0 *vs* 30 min post administration). Note: neuroglucopenia-induced hyperglycemia is the typical readout of the glucose counterregulatory response (CRR). (**C, D**) Glucose dynamics during the oral glucose tolerance test (OGTT, **C**) and the insulin tolerance test (ITT, **D**). No differences were observed between groups. (**E**) Food intake in overnight fasted Sham^PVT^ and 6-OHDA^PVT^ mice (refeeding). (**F**) Water intake in overnight water-deprived Sham^PVT^ and 6-OHDA^PVT^ mice. (**G**) Food intake in overnight water-deprived Sham^PVT^ and 6-OHDA^PVT^ mice. (**H**) Food intake in Sham^PVT^ and 6-OHDA^PVT^ mice administered with ghrelin (0.5 mg/kg, i.p., orexigenic response). Statistics: * p<0.05, ** p<0.01, *** p<0.001, 6-OHDA^PVT^ *vs* Sham^PVT^ mice. Data are presented as mean ± SD. For statistical details see **Statistical Summary Table**.

Second, we performed a fasting-refeeding test to mimic a negative energy balance (food deprivation). In line with the NSF test (**Figure 5C, D**), after an overnight fasting and a similar loss of body weight (**Suppl. Figure 4C**, https://doi.org/10.6084/m9.figshare.19683228.v1), both experimental groups showed an enhanced food intake with 6-OHDA^PVT^ mice consuming more food than Sham^PVT^ mice (**Figure 6E**). Third, we also decided to measure drinking and food intake in overnight water-deprived animals. Both groups showed again a similar decrease in body weight (**Suppl. Figure 4D**, https://doi.org/10.6084/m9.figshare.19683228.v1). After deprivation, mice were exposed to water. While no differences were observed in drinking behavior (**Figure 6F**), 6-OHDA^PVT^ mice again consumed more food than Sham^PVT^ mice (**Figure 6G**). These results suggest that hindbrain^TH^→PVT inputs scale food intake also following physiological and metabolic stressors.

In order to assess whether orexigenic signals without metabolic challenges also required an intact PVT catecholaminergic transmission, we administered ghrelin in fed mice. As shown in **Figure 6H**, ghrelin similarly induced an increase in food intake in both groups, thereby indicating that canonical orexinergic circuits are not altered.

### Reduced catecholaminergic transmission in the PVT promotes the activation of hypothalamic regions

The above-mentioned results point to hindbrain^TH^→PVT afferents as major actors in scaling food intake and metabolic efficiency. Since these homeostatic functions highly depend on the hypothalamus, classically described as the master regulator of energy balance (Dietrich & Horvath, 2013; Timper & Brüning, 2017), we decided to study whether 6-OHDA^PVT^ mice were characterized by an altered basal activity (cFos-positive cells) of key hypothalamic regions such as the dorsomedial hypothalamus (DMH), the ventromedial hypothalamus (VMH), the lateral hypothalamus (LH) and the arcuate nucleus (Arc). As for **Figure 1**, mice were perfused 1h before the onset of the dark phase. Interestingly, in 6-OHDA^PVT^ mice we observed an increase in cFos-positive cells specifically in the DMH and LH (**Figure 7A-C**) compared to Sham^PVT^ mice, whereas no differences were detected in the VMH and Arc (**Figure 7A, D, E**).

**Figure 7:**
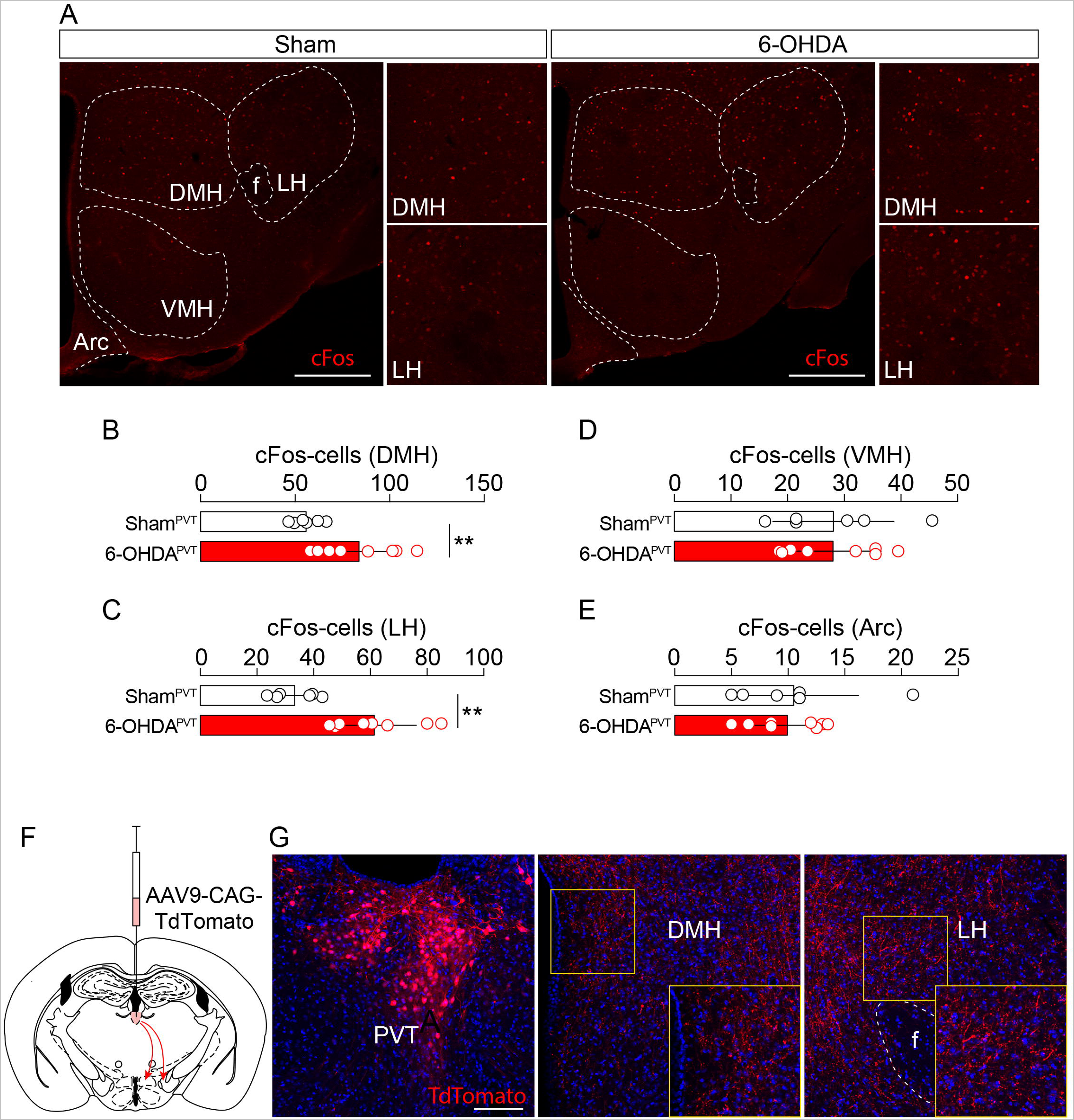
Deletion of hindbrain^TH^→PVT inputs increases cFos expression in the hypothalamus. (**A**) Immunofluorescence detection of cFos in hypothalamic regions of Sham^PVT^ and 6-OHDA^PVT^ mice, notably the dorsomedial hypothalamus (DMH), the lateral hypothalamus (LH), the ventromedial hypothalamus (VMH) and the arcuate nucleus (Arc). Regions of interest are delineated by white dotted lines. Insets represent higher magnifications of DMH and LH regions. Scale bars: 500 µm. Quantification of cFos-positive cells in the DMH (**B**), LH (**C**), VMH (**D**) and Arc (**E**) regions. Statistics: ** p<0.01, 6-OHDA^PVT^ *vs* Sham^PVT^ mice. Abbreviations: f (fornix). (**F**) Drawing indicates the microinjection of AAV9-CAG-TdTomato in the PVT. (**G**) Immunofluorescence detection of TdTomato in the PVT (injection site), DMH and LH (projections). Scale bars: 150 µm. Data are presented as mean ± SD. For statistical details see **Statistical Summary Table**.

In order to see whether PVT-neurons may potentially modulate hypothalamic functions by directly projecting to the DMH and LH, we microinjected an AAV9-CAG-TdTomato virus in the PVT (**Figure 7F, G**). As shown in **Figure 7G**, we observed direct PVT→DMH and PVT→LH projections. These results suggest that the increased activity of PVT-neurons following catecholamines depletion (**Figure 1**) may impact, directly (**Figure 7G**) and/or indirectly (polysynaptic circuits), on the regulatory activity of the hypothalamus which may ultimately result in the modulation of food intake and metabolic efficiency.

## Discussion

The regulation of food intake represents one of the most complex biological functions. Pivotal for adaptation and survival, the regulatory processes underlying feeding are constantly shaped by signals reflecting/sensing physiological adjustments. In virtue of their heterogeneity (exteroceptive *vs* interoceptive sources), stress-like allostatic stimuli are indeed powerful modulators of food intake, feeding habits and metabolic adaptations.

In this study, we report that hindbrain catecholaminergic (putative noradrenergic) inputs to the PVT play a key role in modulating food intake and metabolic efficiency in stress-related contexts. In fact, permanent ablation of TH-afferents to the PVT resulted in enhanced food intake, adjusted metabolic efficiency and nutrient partitioning whenever environmental, behavioral, physiological and/or metabolic (acute/transient) stressors were introduced as dependent variables of feeding behaviors. In particular, 6-OHDA^PVT^ mice were resistant or less sensitive to environmental and behavioral stress-induced hypophagia and showed enhanced feeding patterns following physiological and metabolic challenges. The different nature of stressors used in this study highlights the highly conserved role of CA inputs to the PVT in readily scaling feeding and metabolic adaptations. Moreover, beyond the impact on feeding and metabolic efficiency, it is important to mention that stressors-elicited homeostatic adaptations such as energy expenditure and thermogenesis did not depend on PVT CA inputs, suggesting a functional tropism of hindbrain^TH^→PVT circuits toward food intake on one hand and peripheral nutrient partitioning on the other. This is relevant since previous studies have suggested that PVT-neurons, by facilitating hypothalamic-pituitary-adrenal (HPA) responses (Bhatnagar *et al*., 2000), may contribute to the regulation of core temperature rhythms as well as body weight gain in chronically stressed rats (Bhatnagar & Dallman, 1999). Whether distinct PVT networks (inputs/outputs) are differently engaged by acute *vs* chronic stressors on the regulation of body hemostasis will deserve in-depth investigations. Overall, these results indicate that the PVT, a key node of the limbic circuitry (Barson *et al*., 2020), contributes to the elaboration of food-related decisions and strategies by integrating, among others, also catecholaminergic information. In addition, our results provide new evidence for the existence of distinct, but converging, hindbrain CA inputs (a subset of NTS^TH^- and LC^TH^-neurons) capable of gating food-related homeostatic adaptations under transient stress-like allostatic stimuli.

Surprisingly, we found that the NTS represented one of the major sources of TH-positive projections to the PVT. In fact, local microinjection of 6-OHDA resulted in a significant reduction of TH-expressing neurons in the NTS and to a lesser extend in the LC, sparing CA neurons in the medulla (A1 area) and the hypothalamus, the latter known to send only minor scattered projections to the PVT (Wang *et al*., 2021b). Although to our knowledge no other studies have functionally assessed this NTS^TH^→PVT connection, the impact of 6-OHDA^PVT^ on NTS^TH^-neurons is in line with the presence of a dense plexus of PVT-reaching TH fibers when fluorescent recombinant markers are directly microinjected in the NTS of *Th*-Cre animals (Aklan *et al*., 2020) as well as with a recent retrograde viral study identifying a subset of NTS^TH^-neurons projecting to the PVT (Kirouac *et al*., 2022). This evidence is of paramount important since NTS and LC catecholaminergic neurons, by converging onto the PVT, may synergistically modulate feeding patterns and metabolic efficiency in stressogenic contexts. In fact, NTS^TH^- and LC^TH^-neurons are well-known to modulate food intake and stress/novelty, respectively (McCall *et al*., 2015; Roman *et al*., 2016; Takeuchi *et al*., 2016).

Recent studies have shown that activation of CA-releasing NTS^DBH/TH^-neurons may result in a reduction (Roman *et al*., 2016; Chen *et al*., 2020) as well as in an increase (Aklan *et al*., 2020; Chen *et al*., 2020) of food intake depending on CA cell types and/or projection sites [parabrachial nucleus (PBN) *vs* arcuate nucleus (Arc)]. Although not directly assessed in our study, our results suggest that NTS^TH^→PVT projecting neurons may serve as anorexigenic stimuli since their ablation enhances food intake under stress-related contexts. Indeed, it may be legitimately argued that the use of local microinjections of 6-OHDA may result in the degeneration of LC^TH^- and NTS^TH^-neurons projecting to the PVT but eventually to also other brain regions. However, loss of hindbrain^TH^ neurons projecting to the medial hypothalamus resulted in a loss of glucoprivation-induced feeding (Fraley & Ritter, 2003; Hudson & Ritter, 2004), while in our study we show an increased food consumption under glucoprivic conditions when hindbrain^TH^→PVT projections were ablated. Moreover, opto-activation of NTS^TH^→Arc projections leads to an increase in food intake (Aklan *et al*., 2020), whereas in our case enhanced food intake was elicited by the absence of hindbrain^TH^→PVT projections. These effects may substantiate the functional selectivity of hindbrain^TH^→PVT projections in the adaptive responses of feeding, metabolic efficiency and nutrient partitioning to stress-related contexts. Indeed, future investigations using projection-specific optogenetics and/or chemogenetics may definitely help in dissecting out the distinct and, most importantly, synergistic roles of NTS^TH^→PVT and LC^TH^→PVT transmissions in guiding the tight interaction between food intake, metabolism and stress-like allostatic stimuli.

Moreover, we observed that deletion of hindbrain^TH^→PVT inputs led to an increase in PVT-cells activity (cFos), suggesting that catecholamines may act, directly and/or indirectly, as negative modulators onto PVT-neurons. At first, this is surprising and counterintuitive since *ex vivo* bath-applications of DA (precursor of NE) or NE, which can be synaptically co-released from CA terminals (Smith & Greene, 2012; Kempadoo *et al*., 2016), lead to disinhibition [DA, (Beas *et al*., 2018)] or activation [NE, (Wang *et al*., 2021b)] of PVT-neurons. However, it should be mentioned that mid-posterior PVT-neurons express DA D2 and D3 receptors (Rieck *et al*., 2004; Clark *et al*., 2017; Beas *et al*., 2018; Gao *et al*., 2020) as well as NE receptors such as the α1, α2, β1 and β2 receptors (Rainbow *et al*., 1984; Pieribone *et al*., 1994; Rosin *et al*., 1996). Indeed, (*i*) how this variety of G-protein-coupled receptors (G_i_-, G_s_-, G_q_- and β-arrestin-coupled receptors) mechanistically contribute to the overall modulation of PVT-neurons and (*ii*) whether 6-OHDA-induced TH deletion reorganizes the expression of the above-mentioned CA receptors in the PVT require future investigations. Although our results cannot distinguish between the functional roles of distinct catecholamines onto their associated multiple receptors located onto PVT-neurons, it is worth to mention that the PVT does not receive pure DA-fibers from the midbrain (SNc and VTA) (Li *et al*., 2014; Papathanou *et al*., 2019) and that direct 6-OHDA-induced LC^TH^-neurons loss was associated to an increased expression of Gq-coupled α1 receptor in the thalamus (Szot *et al*., 2012a) which may explain, at least in part, the increase in PVT cFos-neurons.

The enhanced activation of PVT-neurons in 6-OHDA^PVT^ mice and the associated feeding behaviors are in line with reports showing that stressors as well as hunger are able to activate PVT-neurons (Bubser & Deutch, 1999; Beas *et al*., 2018; Hua *et al*., 2018). In addition, our results are also in line with a recent report showing that activation of PVT-neurons by oxytocin was efficient in suppressing stress-induced hypophagia (Barrett *et al*., 2021). However, it is worth to mention that satietogenic signals are also able to activate PVT-neurons (Ong *et al*., 2017), therefore indicating that PVT excitatory (glutamate) neurons may be actually segregated into several cell types with distinct neurochemical, cellular and functional features. This is already supported by the existence of at least two neuronal populations [galanin- and dopamine 2 receptor (D2R)-positive neurons, (Gao *et al*., 2020)] and, as already suggested by the presence of several neuropeptides in PVT-neurons (Curtis *et al*., 2020), it may not be hazardous to hypothesize that future cell type-specific transcriptomic analyses will reveal new sub-families and clusters.

The homeostatic processes underlying food intake, energy balance and metabolic efficiency strongly depend on the activity of the hypothalamus (Dietrich & Horvath, 2013; Timper & Brüning, 2017). We observed that depletion of TH-afferents to the PVT resulted not only in the activation of PVT-neurons but also in the concomitant activation of hypothalamic regions, notably the lateral (LH) and the dorsomedial (DMH) hypothalamus. Indeed, activation of LH and DMH cell types has been shown to promote feeding (Jennings *et al*., 2015; Navarro *et al*., 2016; Otgon-Uul *et al*., 2016; Jeong *et al*., 2017) even following anxiogenic environmental cues (Cassidy *et al*., 2019). Although we cannot rule out yet whether and how the adaptive activation of PVT-neurons following TH deletion may be responsible for the direct activation of LH and DMH regions, it is interesting to note that PVT excitatory (glutamate) neurons also project to the hypothalamus (Engelke *et al*., 2021; Li *et al*., 2021), therefore potentially modulating feeding and energy homeostasis. This is also supported by our viral tracing strategy which revealed direct PVT→DMH/LH projections. However, we cannot formally exclude that the partial loss of hindbrain TH-neurons may impact on the hypothalamic activity in virtue of other circuits (hindbrain^TH^→hypothalamus and/or hindbrain^TH^→PBN→hypothalamus paths). Indeed, while the existence of a hypothalamus→PVT→accumbal circuit seems critical for behavioral adaptations (Betley *et al*., 2013; Zhang & van den Pol, 2017; Otis *et al*., 2019; Meffre *et al*., 2019; Zhang *et al*., 2020; Iglesias & Flagel, 2021; Engelke *et al*., 2021), our results, together with previous and recent literature (Otake *et al*., 1994; Ong *et al*., 2017; Beas *et al*., 2018; Sofia Beas *et al*., 2020; Li *et al*., 2021), also suggest a hindbrain→PVT→hypothalamus path that may regulate homeostatic functions requiring the integration of exteroceptive and interoceptive signals.

In conclusion, the PVT has been classically positioned as a functional node of the limbic circuit (Barson *et al*., 2020). Only recently the hypothesis of the PVT as a homeostatic relay has been proposed (Penzo & Gao, 2021). Altogether, our results support the working hypothesis according to which the PVT, through its afferent connections with NTS^TH^- and LC^TH^-neurons, may represent a functional interface between homeostatic and emotional states, thereby leading to allostatic adaptations. This study, besides highlighting the existence of a dual hindbrain-to-thalamus connection, (*i*) provides new evidence to better understand the dynamic processes underlying the regulation of food intake and energy metabolism, and (*ii*) may serve as starting step to explore the functional relationships and comorbidities between psychiatric (stress) and metabolic (anorexia, obesity, binge eating) disorders.

## Supporting information

Statistical Summary Table

## Acknowledgments

We thank Olja Kacanski for administrative support; Isabelle Le Parco, Ludovic Maingault, Angélique Dauvin, Aurélie Djemat, Florianne Michel and Daniel Quintas for animals’ care; Benoit Bertrand for technical help. We acknowledge the *Functional and Physiological Exploration platform* (FPE) of the Université de Paris (BFA, UMR 8251) and the animal facility Buffon of the Institut Jacques Monod. This work was supported by the Fyssen Foundation, Nutricia Research Foundation, Allen Foundation Inc., *Agence Nationale de la Recherche* (ANR-21-CE14-0021-01), *Fédération pour la Recherche sur le Cerveau* and *Association France Parkinson*, Université Paris Cité and CNRS. G.L. was supported by the China Scholarship Council (CSC) fellowship.

## Author Contributions

C.D. performed and analyzed most of the experiments. G.L. performed immunofluorescence studies. J.C. performed surgeries. S.L. provided critical feedback. G.G. conceived and supervised the whole project, and wrote the manuscript with contribution from all coauthors.

## Data availability statement

All data are presented in the manuscript or supplementary information. For Suppl. Figures see https://doi.org/10.6084/m9.figshare.19683228.v1.

## Figure legends

**Suppl. Figure 1: Detection of DAT-positive fibers in the PVT and TH-positive neurons in the hypothalamus following PVT 6-OHDA microinjections.** (**A**) Immunofluorescence detection of the dopamine transporter (DAT, red) and DAPI (blue) in the PVT, tail of the striatum (TS) and central amygdala (CeA). The lack of DAT-positive fibers in the PVT suggests no direct projections from dopamine-(DAT)-containing midbrain regions. Scale bar: 150 µm. (**B**) Immunofluorescence detection of TH (red) and DAPI (blue) in the hypothalamus of Sham^PVT^ and 6-OHDA^PVT^ mice. Scale bar: 500 µm.

**Suppl. Figure 2: Hindbrain^TH^→PVT inputs dos not alter palatability for HFHS diet.** (**A**) Food intake in Sham^PVT^ and 6-OHDA^PVT^ mice following time-locked feeding (1h) of high-fat high-sugar (HFHS) diet during two consecutive days. Statistics: ** p<0.01, Day2 *vs* Day1 (Sham^PVT^ mice); ^##^ p<0.01, Day2 *vs* Day1 (6-OHDA^PVT^ mice). No differences between experimental groups. Data are presented as mean ± SD. For statistical details see **Statistical Summary Table**.

**Suppl. Figure 3: Confirmation of sensitivity to novelty-induced hypophagia in Sham^PVT^ and 6-OHDA^PVT^ mice before the inverted cycle.** (**A**) Cumulative food intake during the dark phase (spontaneous eating) in Sham^PVT^ and 6-OHDA^PVT^ mice during the first exposure to a novel environment (calorimetric chambers). (**B**) Total food intake during the dark phase. Statistics: *** p<0.001, ** p<0.01, 6-OHDA^PVT^ *vs* Sham^PVT^ mice. Data are presented as mean ± SD. For statistical details see **Statistical Summary Table**.

**Suppl. Figure 4: Effect of fasting and water deprivation in Sham^PVT^ and 6-OHDA^PVT^ mice.** (**A**) Loss of body weight and (**B**) glucose variations in overnight fasted animals used for the novelty-suppressed feeding (NSF) test. (**C**) Loss of body weight in overnight fasted animals before the refeeding schedule. (**D**) Loss of body weight in overnight water-deprived animals before having access to water and chow pellets. No differences between experimental groups. Data are presented as mean ± SD. For statistical details see **Statistical Summary Table**.

