## Supplementary figures and images for "Hindbrain catecholaminergic inputs to the paraventricular thalamus scale feeding and metabolic efficiency in stress-related contexts"

### Suppl. Figure 1

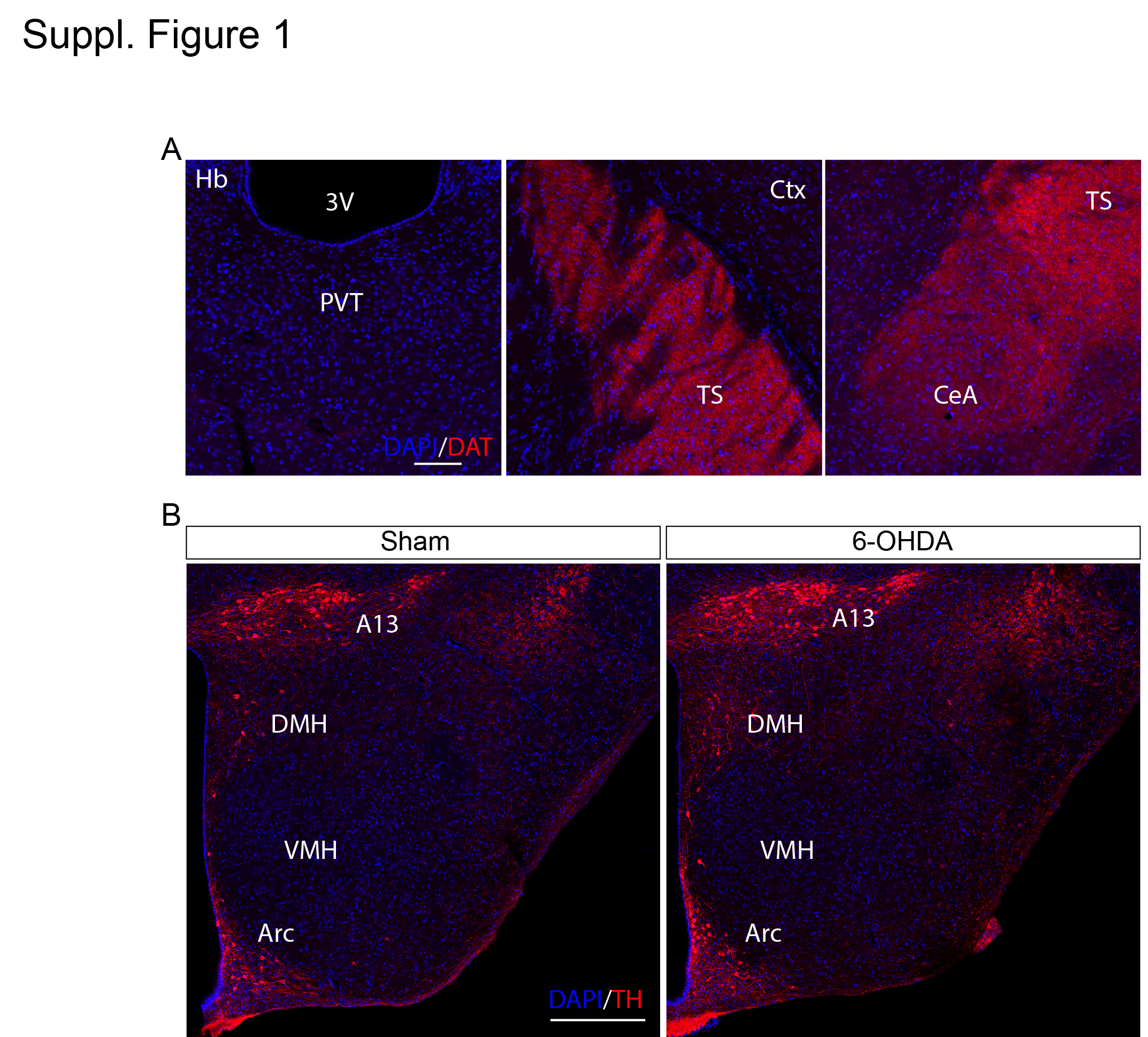

### Suppl. Figure 2

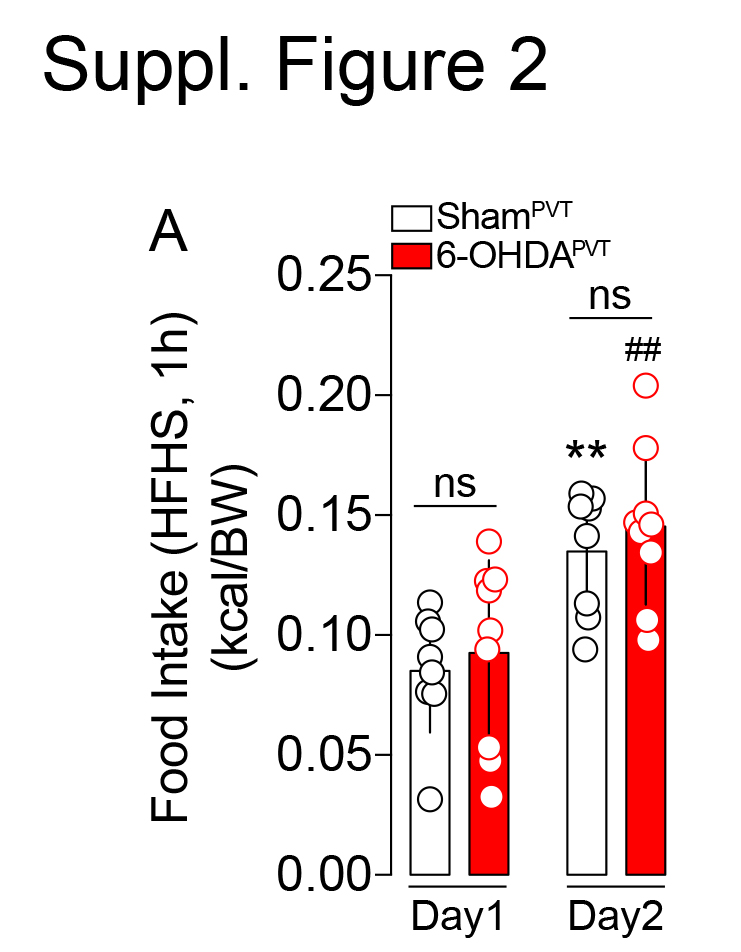

### Suppl. Figure 3

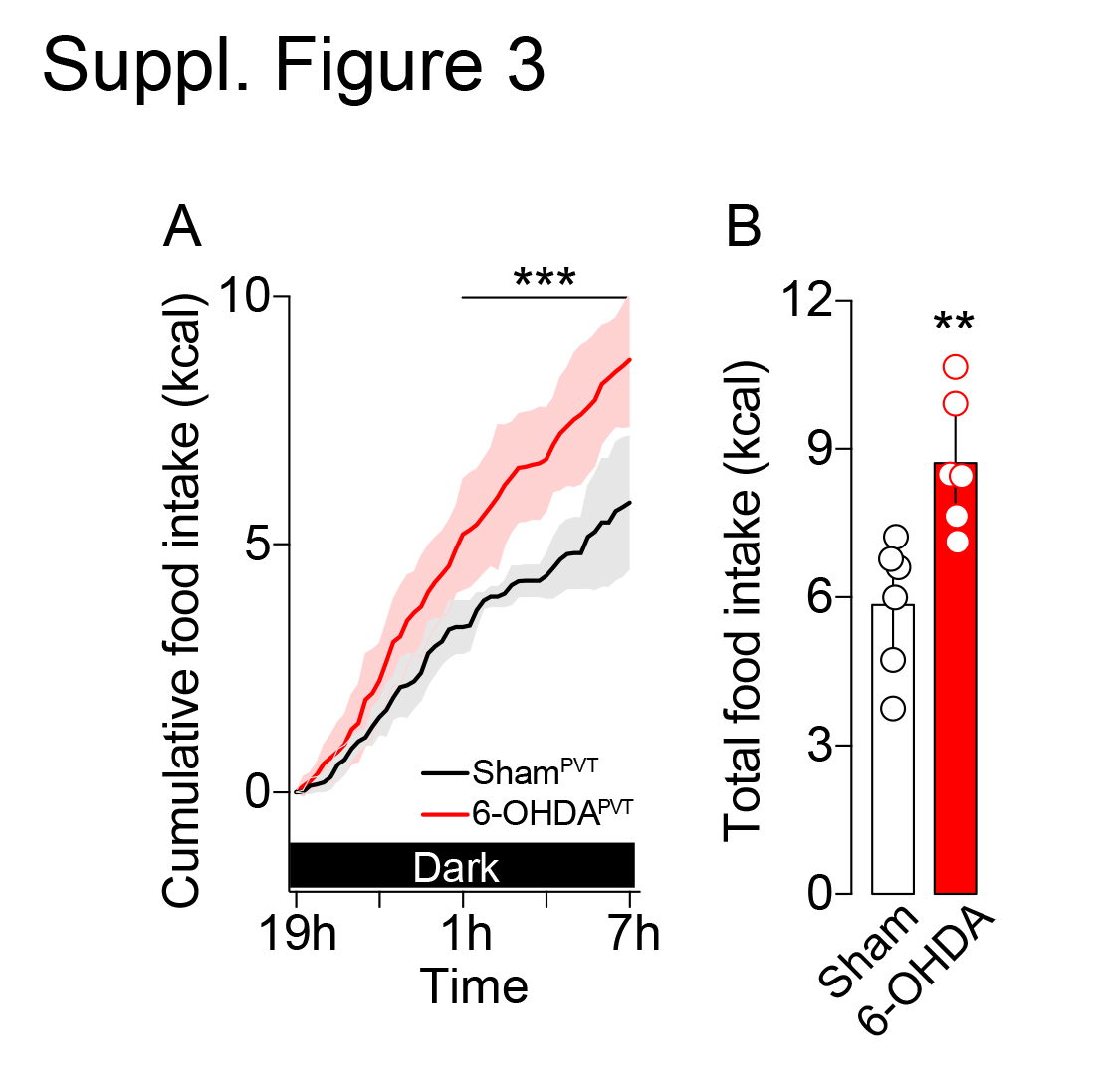

### Suppl. Figure 4

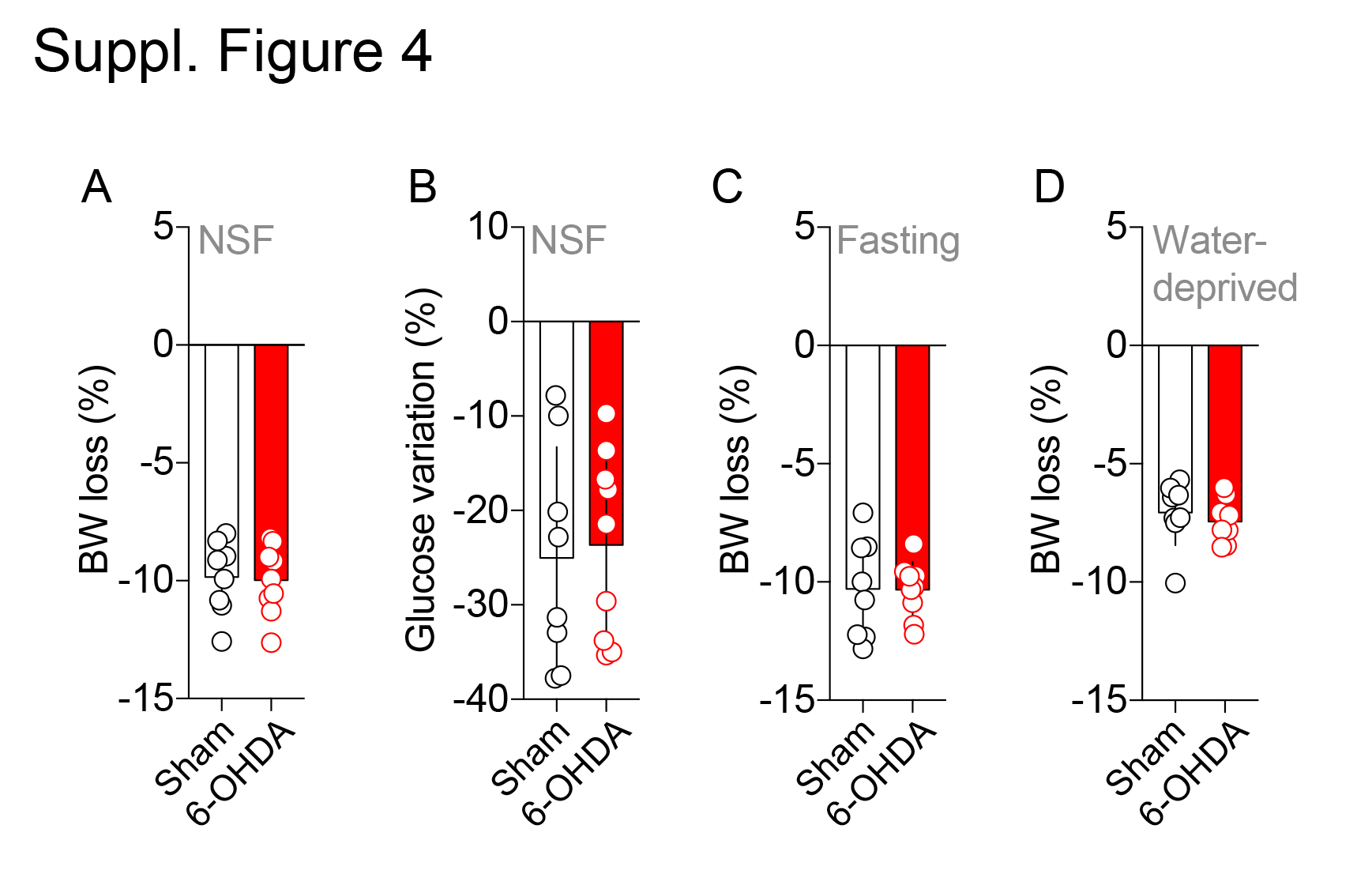
