## Supplementary material for "Hindbrain catecholaminergic inputs to the paraventricular thalamus scale feeding and metabolic efficiency in stress-related contexts": Statistical Summary Table

| **Statistics of Figure 1** | | | | | |
| --- | --- | --- | --- | --- | --- |
| **Figure panels** | | **n** | **Statistical analysis** | **F-value** | **p-value** |
| Fig 1D | TH-neurons in NTS | Sham^PVT^ n=6,  6-OHDA^PVT^ n=8 | Unpaired t-test |  | p = 0.0068,  t=3.259, df=12 |
|  | TH-neurons in area A1 | Sham^PVT^ n=6,  6-OHDA^PVT^ n=8 | Unpaired t-test |  | p = 0.9217,  t=0.100, df=12 |
|  | TH-neurons in LC | Sham^PVT^ n=6,  6-OHDA^PVT^ n=8 | Unpaired t-test |  | p = 0.0359,  t=2.363, df=12 |
| Fig 1F | cFos-neurons in PVT | Sham^PVT^ n=6,  6-OHDA^PVT^ n=8 | Unpaired t-test |  | p = 0.0003,  t=5.016, df=12 |

| **Statistics of Figure 2** | | | | | | |
| --- | --- | --- | --- | --- | --- | --- |
| **Figure panels** | | **n** | **Statistical analysis** | | **F-value** | **p-value** |
| Fig 2A | Body weight (g) | Sham^PVT^ n=8,  6-OHDA^PVT^ n=10 | Unpaired t-test | |  | p = 0.8800 |
| Fig 2C | Food intake in single-housed mice (D1 to D3) | Sham^PVT^ n=8,  6-OHDA^PVT^ n=10 | 2-way ANOVA | interaction | F_(2, 32)_ = 3.466 | p = 0.0434 |
|  |  |  |  | time | F_(2, 32)_ = 35.12 | p < 0.0001 |
|  |  |  |  | groups | F_(1, 16)_ = 9.709 | p = 0.0067 |
| Fig 2E | Food intake during exposure to novel environment | Sham^PVT^ n=8,  6-OHDA^PVT^ n=10 | 2-way ANOVA | interaction | F_(144, 1536)_ = 4.16 | p < 0.0001 |
|  |  |  |  | time | F_(48, 1536)_ = 583.8 | p < 0.0001 |
|  |  |  |  | groups | F_(3, 32)_ = 6.342 | p = 0.0017 |
| Fig 2E^1^ | Food intake during exposure to novel environment | Sham^PVT^ n=8,  6-OHDA^PVT^ n=10 | 2-way ANOVA | interaction | F_(1, 16)_ = 9.023 | p = 0.0084 |
|  |  |  |  | time | F_(1, 16)_ = 26.62 | p < 0.0001 |
|  |  |  |  | groups | F_(1, 16)_ = 2.996 | p = 0.1027 |
| Fig 2F | Locomotor activity during exposure to a novel environment | Sham^PVT^ n=8,  6-OHDA^PVT^ n=10 | 2-way ANOVA | interaction | F_(144, 1536)_ = 1.61 | p < 0.0001 |
|  |  |  |  | time | F_(48, 1536)_ = 13.89 | p < 0.0001 |
|  |  |  |  | groups | F_(3, 32)_ = 13.12 | p < 0.0001 |
| Fig 2F^1^ | Locomotor activity during exposure to a novel environment | Sham^PVT^ n=8,  6-OHDA^PVT^ n=10 | 2-way ANOVA | interaction | F_(1, 16)_ = 0.0001 | p = 0.9913 |
|  |  |  |  | time | F_(1, 16)_ = 35.55 | p < 0.0001 |
|  |  |  |  | groups | F_(1, 16)_ = 0.0887 | p = 0.7697 |
| Fig 2G | RER during first exposure to novel environment | Sham^PVT^ n=8,  6-OHDA^PVT^ n=10 | 2-way ANOVA | interaction | F_(48, 768)_ = 1.012 | p = 0.4536 |
|  |  |  |  | time | F_(48, 768)_ = 8.807 | p < 0.0001 |
|  |  |  |  | groups | F_(1, 16)_ = 11.49 | p = 0.0037 |
| Fig 2G^1^ | RER during first exposure to novel environment | Sham^PVT^ n=8,  6-OHDA^PVT^ n=10 | Unpaired t-test | |  | p = 0.0037  t=3.389, df=16 |
| Fig 2H | FAO during first exposure to novel environment | Sham^PVT^ n=8,  6-OHDA^PVT^ n=10 | 2-way ANOVA | interaction | F_(48, 768)_ = 0.696 | p = 0.9417 |
|  |  |  |  | time | F_(48, 768)_ = 14.05 | p < 0.0001 |
|  |  |  |  | groups | F_(1, 16)_ = 36.42 | p = 0.0026 |
| Fig 2H^1^ | FAO during first exposure to novel environment | Sham^PVT^ n=8,  6-OHDA^PVT^ n=10 | Unpaired t-test | |  | p = 0.0026  t=3.558, df=16 |
| Fig 2I | EE during first exposure to novel environment | Sham^PVT^ n=8,  6-OHDA^PVT^ n=10 | 2-way ANOVA | interaction | F_(48, 768)_ = 0.806 | p = 0.8238 |
|  |  |  |  | time | F_(48, 768)_ = 6.441 | p < 0.0001 |
|  |  |  |  | groups | F_(1, 16)_ = 0.282 | p = 0.6027 |
| Fig 2I^1^ | EE during first exposure to novel environment | Sham^PVT^ n=8,  6-OHDA^PVT^ n=10 | Unpaired t-test | |  | p = 0.6449  t=0.4697, df=16 |

| **Statistics of Figure 3** | | | | | | |
| --- | --- | --- | --- | --- | --- | --- |
| **Figure panels** | | **n** | **Statistical analysis** | | **F-value** | **p-value** |
| Fig 3B | Total distance | Sham^PVT^ n=8,  6-OHDA^PVT^ n=9 | Unpaired t-test | |  | p = 0.8083  t=0.247, df=15 |
| Fig 3C | Entries in center zone | Sham^PVT^ n=8,  6-OHDA^PVT^ n=9 | Unpaired t-test | |  | p = 0.1568  t=1.491, df=15 |
| Fig 3D | % of time in center zone | Sham^PVT^ n=8,  6-OHDA^PVT^ n=9 | Unpaired t-test | |  | p = 0.0653  t=1.989, df=15 |
| Fig 3E | Mean exploration visits (s) | Sham^PVT^ n=8,  6-OHDA^PVT^ n=9 | Unpaired t-test | |  | p = 0.7140  t=0.3735, df=15 |
| Fig 3F | Distance in center (cm) | Sham^PVT^ n=8,  6-OHDA^PVT^ n=9 | Unpaired t-test | |  | p = 0.1186  t=1.655, df=15 |
| Fig 3H | BAT temperature  (before and after OF) | Sham^PVT^ n=8,  6-OHDA^PVT^ n=9 | 2-way ANOVA | interaction | F_(1, 15)_ = 0.0007 | p = 0.9934 |
|  |  |  |  | time | F_(1, 15)_ = 219.6 | p < 0.0001 |
|  |  |  |  | groups | F_(1, 15)_ = 0.401 | p = 0.5361 |
| Fig 3I | Lower back temperature  (before and after OF) | Sham^PVT^ n=8,  6-OHDA^PVT^ n=9 | 2-way ANOVA | interaction | F_(1, 15)_ = 0.0829 | p = 0.7773 |
|  |  |  |  | time | F_(1, 15)_ = 888.5 | p < 0.0001 |
|  |  |  |  | groups | F_(1, 15)_ = 0.3297 | p = 0.5744 |
| Fig 3J | Tail temperature  (before and after OF) | Sham^PVT^ n=8,  6-OHDA^PVT^ n=9 | 2-way ANOVA | interaction | F_(1, 15)_ = 3.412 | p = 0.1413 |
|  |  |  |  | time | F_(1, 15)_ = 1840 | p < 0.0001 |
|  |  |  |  | groups | F_(1, 15)_ = 3.204 | p = 0.0936 |

| **Statistics of Figure 4** | | | | | | |
| --- | --- | --- | --- | --- | --- | --- |
| **Figure panels** | | **n** | **Statistical analysis** | | **F-value** | **p-value** |
| Fig 4B | Food intake during standard and inverted cycles  (Sham^PVT^ mice) | Sham^PVT^  n=6 | 2-way ANOVA | interaction | F_(96, 960)_ = 15.24 | p < 0.0001 |
|  |  |  |  | time | F_(96, 960)_ = 385.4 | p < 0.0001 |
|  |  |  |  | groups | F_(1, 10)_ = 1.324 | p = 0.2766 |
| Fig 4B^1^ | Food intake during standard and inverted cycles  (6-OHDA^PVT^ mice) | 6-OHDA^PVT^ n=6 | 2-way ANOVA | interaction | F_(96, 960)_ = 18.13 | p < 0.0001 |
|  |  |  |  | time | F_(96, 960)_ = 394.9 | p < 0.0001 |
|  |  |  |  | groups | F_(1, 10)_ = 0.0884 | p = 0.7723 |
| Fig 4C | Food intake during standard and inverted cycles | Sham^PVT^ n=6,  6-OHDA^PVT^ n=6 | 2-way ANOVA | interaction | F_(1, 10)_ = 7.518 | p = 0.0208 |
|  |  |  |  | time | F_(1, 10)_ = 1.991 | p = 0.1886 |
|  |  |  |  | groups | F_(1, 10)_ = 0.1222 | p = 0.7339 |
| Fig 4D | EE during standard and inverted cycles  (Sham^PVT^ mice) | Sham^PVT^  n=6 | 2-way ANOVA | interaction | F_(96, 960)_ = 13.4 | p < 0.0001 |
|  |  |  |  | time | F_(96, 960)_ = 9.439 | p < 0.0001 |
|  |  |  |  | groups | F_(1, 10)_ = 0.0252 | p = 0.8770 |
| Fig 4D^1^ | EE during standard and inverted cycles  (6-OHDA^PVT^ mice) | 6-OHDA^PVT^ n=6 | 2-way ANOVA | interaction | F_(96, 960)_ = 13.12 | p < 0.0001 |
|  |  |  |  | time | F_(96, 960)_ = 9.672 | p < 0.0001 |
|  |  |  |  | groups | F_(1, 10)_ = 0.1402 | p = 0.7159 |
| Fig 4E | EE during standard and inverted cycles  (7h00-19h00) | Sham^PVT^ n=6,  6-OHDA^PVT^ n=6 | 2-way ANOVA | interaction | F_(1, 10)_ = 1.28 | p = 0.2843 |
|  |  |  |  | time | F_(1, 10)_ = 85.99 | p < 0.0001 |
|  |  |  |  | groups | F_(1, 10)_ = 0.611 | p = 0.4525 |
| Fig 4E^1^ | EE during standard and inverted cycles  (19h00-7h00) | Sham^PVT^ n=6,  6-OHDA^PVT^ n=6 | 2-way ANOVA | interaction | F_(1, 10)_ = 0.2259 | p = 0.6448 |
|  |  |  |  | time | F_(1, 10)_ = 201.4 | p < 0.0001 |
|  |  |  |  | groups | F_(1, 10)_ = 0.6147 | p = 0.4512 |
| Fig 4F | FAO during standard and inverted cycles  (Sham^PVT^ mice) | Sham^PVT^  n=6 | 2-way ANOVA | interaction | F_(96, 960)_ = 17.66 | p < 0.0001 |
|  |  |  |  | time | F_(96, 960)_ = 20.73 | p < 0.0001 |
|  |  |  |  | groups | F_(1, 10)_ = 5.119 | p = 0.0472 |
| Fig 4F^1^ | FAO during standard and inverted cycles  (6-OHDA^PVT^ mice) | 6-OHDA^PVT^ n=6 | 2-way ANOVA | interaction | F_(96, 960)_ = 15.54 | p < 0.0001 |
|  |  |  |  | time | F_(96, 960)_ = 20.33 | p < 0.0001 |
|  |  |  |  | groups | F_(1, 10)_ = 4.577 | p = 0.0681 |
| Fig 4G | FAO during standard and inverted cycles  (7h00-19h00) | Sham^PVT^ n=6,  6-OHDA^PVT^ n=6 | 2-way ANOVA | interaction | F_(1, 10)_ = 14.47 | p = 0.0035 |
|  |  |  |  | time | F_(1, 10)_ = 45.75 | p < 0.0001 |
|  |  |  |  | groups | F_(1, 10)_ = 0.2171 | p = 0.6513 |
| Fig 4G^1^ | FAO during standard and inverted cycles  (19h00-7h00) | Sham^PVT^ n=6,  6-OHDA^PVT^ n=6 | 2-way ANOVA | interaction | F_(1, 10)_ = 0.2233 | p = 0.6467 |
|  |  |  |  | time | F_(1, 10)_ = 40.2 | p < 0.0001 |
|  |  |  |  | groups | F_(1, 10)_ = 0.4982 | p = 0.4964 |
| Fig 4H | RER during standard and inverted cycles  (Sham^PVT^ mice) | Sham^PVT^  n=6 | 2-way ANOVA | interaction | F_(96, 960)_ = 21.83 | p < 0.0001 |
|  |  |  |  | time | F_(96, 960)_ = 21.83 | p < 0.0001 |
|  |  |  |  | groups | F_(1, 10)_ = 3.477 | p = 0.0918 |
| Fig 4H^1^ | RER during standard and inverted cycles  (6-OHDA^PVT^ mice) | 6-OHDA^PVT^ n=6 | 2-way ANOVA | interaction | F_(96, 960)_ = 17.65 | p < 0.0001 |
|  |  |  |  | time | F_(96, 960)_ = 21.21 | p < 0.0001 |
|  |  |  |  | groups | F_(1, 10)_ = 2.464 | p = 0.1475 |
| Fig 4I | RER during standard and inverted cycles  (7h00-19h00) | Sham^PVT^ n=6,  6-OHDA^PVT^ n=6 | 2-way ANOVA | interaction | F_(1, 10)_ = 10.45 | p = 0.0090 |
|  |  |  |  | time | F_(1, 10)_ = 47.92 | p < 0.0001 |
|  |  |  |  | groups | F_(1, 10)_ = 0.0357 | p = 0.8539 |
| Fig 4I^1^ | RER during standard and inverted cycles  (19h00-7h00) | Sham^PVT^ n=6,  6-OHDA^PVT^ n=6 | 2-way ANOVA | interaction | F_(1, 10)_ = 0.0624 | p = 0.8078 |
|  |  |  |  | time | F_(1, 10)_ = 47.28 | p < 0.0001 |
|  |  |  |  | groups | F_(1, 10)_ = 0.4574 | p = 0.5142 |

| **Statistics of Figure 5** | | | | | | |
| --- | --- | --- | --- | --- | --- | --- |
| **Figure panels** | | **n** | **Statistical analysis** | | **F-value** | **p-value** |
| Fig 5B | Latency to first bite (s) during NSF test | Sham^PVT^ n=8,  6-OHDA^PVT^ n=9 | Unpaired t-test | |  | p = 0.5792  t=0.567, df=15 |
| Fig 5C | Cumulative Food Intake (kcal) during NSF test | Sham^PVT^ n=8,  6-OHDA^PVT^ n=9 | Unpaired t-test | |  | p = 0.0096  t=2.969, df=15 |
| Fig 5D | Food Intake (BW/kcal) during NSF test | Sham^PVT^ n=8,  6-OHDA^PVT^ n=9 | Unpaired t-test | |  | p = 0.0106  t=2.92, df=15 |
| Fig 5E | Cumulative Food Intake (kcal) after restraint test | Sham^PVT^ n=8,  6-OHDA^PVT^ n=9 | 2-way ANOVA | interaction | F_(1, 15)_ = 13.66 | p = 0.0022 |
|  |  |  |  | time | F_(1, 15)_ = 92.34 | p < 0.0001 |
|  |  |  |  | groups | F_(1, 15)_ = 6.11 | p = 0.0259 |

| **Statistics of Figure 6** | | | | | | |
| --- | --- | --- | --- | --- | --- | --- |
| **Figure panels** | | **n** | **Statistical analysis** | | **F-value** | **p-value** |
| Fig 6A | 2-DG-induced food intake | Sham^PVT^ n=8,  6-OHDA^PVT^ n=9 | 2-way ANOVA | interaction | F_(3, 45)_ = 2.876 | p = 0.0464 |
|  |  |  |  | time | F_(3, 45)_ = 77.05 | p < 0.0001 |
|  |  |  |  | groups | F_(1, 15)_ = 6.826 | p = 0.0196 |
| Fig 6B | 2-DG-induced counterregulatory glucose reponse | Sham^PVT^ n=8,  6-OHDA^PVT^ n=9 | Unpaired t-test | |  | p = 0.6123  t=0.5176, df=15 |
| Fig 6C | OGTT | Sham^PVT^ n=7,  6-OHDA^PVT^ n=9 | 2-way ANOVA | interaction | F_(6, 84)_ = 1.33 | p = 0.2528 |
|  |  |  |  | time | F_(6, 84)_ = 109.8 | p < 0.0001 |
|  |  |  |  | groups | F_(1, 14)_ = 0.3563 | p = 0.5601 |
| Fig 6D | ITT | Sham^PVT^ n=8,  6-OHDA^PVT^ n=9 | 2-way ANOVA | interaction | F_(7, 105)_ = 4.7 | p = 0.0001 |
|  |  |  |  | time | F_(7,105)_ = 125.9 | p < 0.0001 |
|  |  |  |  | groups | F_(1, 15)_ = 0.5018 | p = 0.4896 |
| Fig 6E | Fasting + Refeeding | Sham^PVT^ n=8,  6-OHDA^PVT^ n=9 | 2-way ANOVA | interaction | F_(3, 45)_ = 2.072 | p = 0.1173 |
|  |  |  |  | time | F_(3,45)_ = 119.6 | p < 0.0001 |
|  |  |  |  | groups | F_(1, 15)_ = 19.96 | p = 0.0005 |
| Fig 6F | Water Intake  (water-deprived mice) | Sham^PVT^ n=8,  6-OHDA^PVT^ n=9 | 2-way ANOVA | interaction | F_(5, 75)_ = 0.8967 | p = 0.4879 |
|  |  |  |  | time | F_(5,75)_ = 175.6 | p < 0.0001 |
|  |  |  |  | groups | F_(1, 15)_ = 0.7289 | p = 0.4067 |
| Fig 6G | Food Intake  (water-deprived mice) | Sham^PVT^ n=8,  6-OHDA^PVT^ n=9 | 2-way ANOVA | interaction | F_(5, 75)_ = 1.681 | p = 0.1496 |
|  |  |  |  | time | F_(5,75)_ = 88.11 | p < 0.0001 |
|  |  |  |  | groups | F_(1, 15)_ = 5.99 | p = 0.0272 |
| Fig 6H | Food Intake  (Ghrelin) | Sham^PVT^ n=8,  6-OHDA^PVT^ n=9 | 2-way ANOVA | interaction | F_(3, 45)_ = 0.3487 | p = 0.7902 |
|  |  |  |  | time | F_(3,45)_ = 35.15 | p < 0.0001 |
|  |  |  |  | groups | F_(1, 15)_ = 0.3942 | p = 0.5395 |

| **Statistics of Figure 7** | | | | | |
| --- | --- | --- | --- | --- | --- |
| **Figure panels** | | **n** | **Statistical analysis** | **F-value** | **p-value** |
| Fig 7A | cFos-neurons in DMH | Sham^PVT^ n=6,  6-OHDA^PVT^ n=8 | Unpaired t-test |  | p = 0.0098  t=3.064, df=12 |
| Fig 7B | cFos-neurons in LH | Sham^PVT^ n=6,  6-OHDA^PVT^ n=8 | Unpaired t-test |  | p = 0.0013  t=4.179, df=12 |
| Fig 7C | cFos-neurons in VMH | Sham^PVT^ n=6,  6-OHDA^PVT^ n=8 | Unpaired t-test |  | p = 0.9873  t=0.0163, df=12 |
| Fig 7D | cFos-neurons in Arc | Sham^PVT^ n=6,  6-OHDA^PVT^ n=8 | Unpaired t-test |  | p = 0.8185  t=0.2345, df=12 |

| **Statistics of Suppl. Figure 2** | | | | | | |
| --- | --- | --- | --- | --- | --- | --- |
| **Figure panels** | | **n** | **Statistical analysis** | | **F-value** | **p-value** |
| SFig 2A | HFHS | Sham^PVT^ n=8,  6-OHDA^PVT^ n=9 | 2-way ANOVA | interaction | F_(1, 15)_ = 0.039 | p = 0.8458 |
|  |  |  |  | time | F_(1, 15)_ = 45.97 | p < 0.0001 |
|  |  |  |  | groups | F_(1, 15)_ = 0.4575 | p = 0.5091 |

| **Statistics of Suppl. Figure 3** | | | | | | |
| --- | --- | --- | --- | --- | --- | --- |
| **Figure panels** | | **n** | **Statistical analysis** | | **F-value** | **p-value** |
| SFig 3A | Cumulative food intake, first exposure before inverted cycle | Sham^PVT^ n=6,  6-OHDA^PVT^ n=6 | 2-way ANOVA | interaction | F_(48, 480)_ = 7.547 | p < 0.0001 |
|  |  |  |  | time | F_(48, 480)_ = 180.4 | p < 0.0001 |
|  |  |  |  | groups | F_(1, 10)_ = 20.64 | p = 0.0011 |
| SFig 3B | Cumulative food intake, first exposure before inverted cycle | Sham^PVT^ n=6,  6-OHDA^PVT^ n=6 | Unpaired t-test | |  | p = 0.0041,  t=3.705, df=10 |

| **Statistics of Suppl. Figure 4** | | | | | |
| --- | --- | --- | --- | --- | --- |
| **Figure panels** | | **n** | **Statistical analysis** | **F-value** | **p-value** |
| SFig 4A | Loss body weight (%)  (fasting, NSF) | Sham^PVT^ n=8,  6-OHDA^PVT^ n=9 | Unpaired t-test |  | p = 0.8553,  t=0.185, df=15 |
| SFig 4B | Glucose variations (%)  (fasting, NSF) | Sham^PVT^ n=8,  6-OHDA^PVT^ n=9 | Unpaired t-test |  | p = 0.7981,  t=0.260, df=15 |
| SFig 4C | Loss body weight (%)  (fasting) | Sham^PVT^ n=8,  6-OHDA^PVT^ n=9 | Unpaired t-test |  | p = 0.9655,  t=0.044, df=15 |
| SFig 4D | Loss body weight (%)  (water deprivation) | Sham^PVT^ n=8,  6-OHDA^PVT^ n=9 | Unpaired t-test |  | p = 0.5079,  t=0.678, df=15 |
